# Structural cell biology by mega-expansion microscopy

**DOI:** 10.64898/2026.08.05.743040

**Authors:** Ignacio Vega-Vásquez, Omar Ignacio García-Martínez, Camila García-Navarrete, Gang Wen, Christian Werner, Patrick Eiring, Jorge A. Toledo, Sushovan Chanda, Clinton Gonsalves, Ali H. Shaib, Gislene Pereira, Silvio O. Rizzoli, Ricardo Benavente, Philip Kollmannsberger, Markus Sauer

## Abstract

Structural characterization of macromolecular assemblies within intact cells remains one of the central challenges in cell biology. While cryo-electron tomography provides unparalleled structural information, its applicability is limited by sample thickness, imaging throughput, and accessibility. Fluorescence microscopy offers molecular specificity and compatibility with intact biological specimens but has so far lacked the spatial resolution required to visualize cellular ultrastructure. Here we introduce Mega-expansion microscopy (Mega-ExM), a fluorescence imaging approach that enables structural visualization of whole cells using conventional confocal microscopes. Mega-ExM combines iterative hydrogel expansion with whole-proteome NHS-dye labeling and post-expansion immunostaining to achieve tunable expansion factors of up to ∼1,500-fold while preserving ultrastructure. At expansion factors of 40-260x, Mega-ExM resolves centrioles, mitochondrial cristae, protein-dense domains within mitochondrial cristae consistent with respiratory-chain supercomplexes, the synaptonemal complex, and nuclear pore complexes (NPCs) with high fidelity. Particle averaging of ∼200x expanded NPCs yields reconstructions with a structural resolution of ∼35 Å, approaching what cryo-electron tomography has achieved for selected protein assemblies. By combining molecular specificity, large imaging volumes, and nanoscale structural resolution on conventional fluorescence microscopes, Mega-ExM establishes a broadly accessible platform for in situ structural biology.

## Main text

Understanding the molecular architecture of cells requires imaging methods that combine nanometer-scale spatial resolution with molecular specificity in intact biological specimens. Recent years have seen dramatic advances in spatial resolution achievable by fluorescence microscopy. Single-molecule localization microscopy methods and their derivatives now reach molecular-scale resolution^1–3^, and, when combined with expansion microscopy (ExM)^4^, can even resolve the shapes and molecular structures of isolated proteins^5,6^. However, visualizing the molecular architecture of intact cells at nanometer resolution remains largely confined to cryo-electron tomography (cryo-ET). Cryo-ET has revolutionized structural cell biology by revealing macromolecular assemblies directly within their native cellular environment. However, cryo-ET remains technically demanding, is fundamentally limited by specimen thickness, and provides little molecular specificity without additional labeling strategies^7,8^.

Fluorescence microscopy offers complementary strengths including high molecular specificity, compatibility with living and thick specimens, and broad accessibility, but its spatial resolution remains fundamentally constrained by both diffraction and labeling density. However, even when localization precision approaches a few nanometers or better in some cases, insufficient labeling density, fluorophore interactions, and steric limitations prevent faithful reconstruction of densely packed molecular assemblies^9–11^. Consequently, visualizing the ultrastructure of intact cells at cryo-ET-like resolution has remained beyond the reach of fluorescence microscopy.

Expansion microscopy (ExM) addresses these limitations by physically separating biomolecules before imaging^4^. Expansion improves epitope accessibility, reduces linkage error and electronic fluorophore interactions, and enables dense labeling of intact cellular structures^12–15^. Furthermore, whole-proteome labeling with NHS-functionalized dyes provides structural context reminiscent of electron microscopy^16,17^. Although recent ExM protocols have improved ultrastructure preservation and enabled imaging at 10-20 nm resolution, expansion factors have remained insufficient to visualize the molecular architecture of intact cells at the level routinely achieved by cryo-ET^18–26^. Recently, an interpenetrating hydrogel architecture has been introduced, allowing thousandfold expansion microscopy (1000ExM) and fluorescence imaging with sub-nanometer precision on single-molecule sensitive conventional microscopes. So far, however, 1000ExM has been limited to resolving adjacent amino acid residues in isolated proteins outside of their physiological context^27^.

Here we introduce Mega-ExM, an iterative expansion strategy that preserves protein integrity while achieving tunable expansion factors of up to ∼250-fold for intact cellular structures and beyond 1,000-fold under optimized conditions. By combining iterative hydrogel expansion with whole-proteome NHS labeling and post-expansion immunostaining, Mega-ExM enables structural imaging of entire cells using conventional confocal microscopes. We demonstrate that Mega-ExM resolves mitochondrial cristae, protein-dense domains within mitochondrial cristae consistent with respiratory-chain supercomplexes, centrioles, the synaptonemal complex, and nuclear pore complexes with a level of structural detail approaching that achieved by cryo-ET for selected protein assemblies.

### Mega-ExM provides tunable expansion factors

Iterative ExM protocols initially relied on sequential embedding of biological samples into swellable but cleavable hydrogels to ensure further expansion after the first expansion round (Supplementary Fig. 1). Recent protocols, however, demonstrated that an expanded uncleavable hydrogel can expand further if a second swellable hydrogel is formed in the space opened by the first expansion step (and stabilized by a non-cleavable neutral gel). However, since these approaches used only ∼4x hydrogels, the final expansion factors were limited to 15-20x^17,21–23,26^. We therefore reasoned that one could achieve higher expansion factors by using (*a*) the non-cleavable TREx protocol that achieves expansion factors of up to 10x in a single round of gelation and expansion^20^, and (*b*) the original stabilizing neutral polyacrylamide hydrogel crosslinked with *N,N’*-(1,2-dihydroxyethylene)bis-acrylamide (DHEBA, an acrylamide-crosslinker with a cleavable amidomethylol bond) (Supplementary Fig. 1)^16^. To examine how Mega-ExM might achieve this additional swelling, we systematically varied the sodium acrylate-to-acrylamide ratio in the TREx recipe while keeping the bis-acrylamide (crosslinker) concentration and the total monomer concentration constant (Supplementary Table 1). Our results indicate that the non-cleavable TREx hydrogel does not reach its full expansion capacity during the first expansion round. Instead, it expands only until the electrostatic repulsion between the negatively charged polymer chains reaches equilibrium with the opposing elastic and osmotic forces within the hydrogel network. The polymer can thus further expand in the second and third expansion round if more charges are introduced in the vicinity through re-embedding (Supplementary Fig. 2). For comparison we tested different expansion protocols including proExM^12^, U-ExM^28^, iU-ExM^21^, and Magnify^29^ as starting point, both, iterated with itself and with TREx with special emphasis on the achieved expansion factors. However, none of the methods came close to the performance of TREx-based Mega-ExM (Supplementary Tables 1-2).

The Mega-ExM protocol starts with fixed cells and crosslinks them with 2% acrylamide (AA) and 1.4% formaldehyde (FA) in phosphate-buffered saline (PBS) for at least 3 h at 37°C, before embedding them in a standard TREx gel recipe^20^. Denaturation follows at 85°C or 95°C in 200 mM sodium dodecyl sulfate (SDS), 200 mM NaCl and 50 mM Tris buffer, for 1.5 h at pH = 6.8 or 1 h at pH = 9.0, respectively, depending on the structure of interest (Fig. 1a, see Methods section)^21^. After the first TREx expansion round (T), aliphatic amino groups are labeled with NHS-functionalized dyes, and decrowded proteins can be immunostained with primary and secondary antibodies with improved epitope accessibility. Subsequently, the TREx gel is re-embedded in a neutral cleavable hydrogel (N) and a second swellable non-cleavable TREx hydrogel (T), resulting in “TNT” expanded samples after neutral gel dissolution in 200 mM NaOH (Fig. 1a-d). Applying another round of NT re-embedding, neutral gel dissolution and expansion of TNT expanded gels results in Mega-ExM (TNTNT) samples (Fig. 1a-e). Immunolabeling after the first (T) or second expansion (TNT) round resulted in similar labeling densities and image qualities (Supplementary Fig. 3). Therefore, we used immunolabeling after the first expansion round in all experiments. The performance of TNT-and Mega-ExM was verified using HeLa, COS-7, and Raji B cells, synaptonemal complexes from mouse spermatocytes, and primary hippocampal neurons (Fig. 1b-e, Fig. 2 and Supplementary Fig. 4). In addition, we demonstrated that Mega-ExM can be used successfully to expand tissue slices (Supplementary Fig. 5).

**Figure 1.**
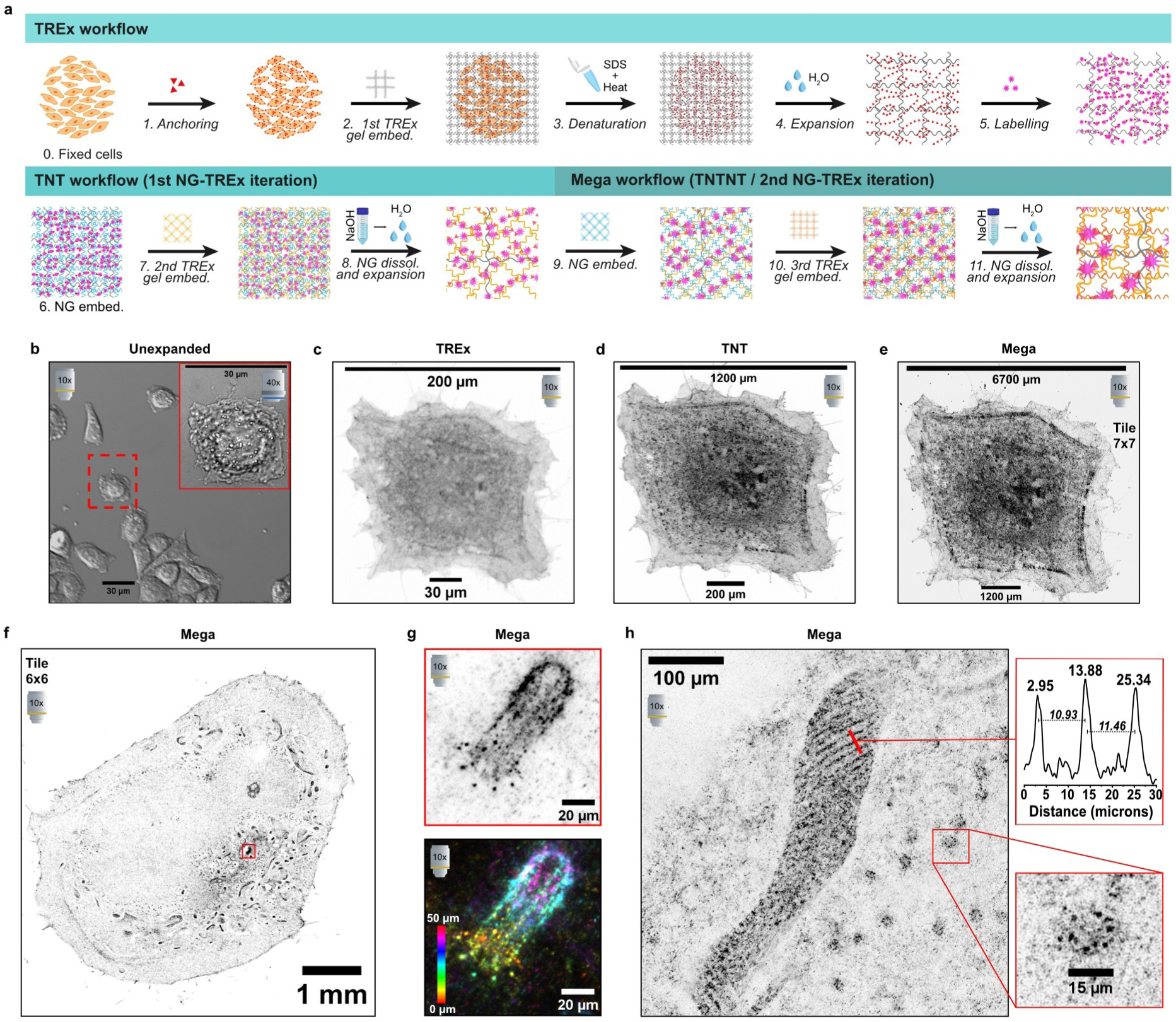
Overview of TNT- and Mega-ExM workflow and output. **a**, Expansion steps including the anchoring, gelation, denaturation, expansion and labeling of a first non-cleavable TREx gel, followed by sequential re-embedding of a cleavable neutral gel and another TREx (this iteration repeated twice in total). **b-e**, The same HeLa cell is shown first unstained and unexpanded in **b**, and then NHS-stained throughout TREx, TNT and Mega steps with approximately ∼7x, 40x and 220x linear expansion factors, respectively, acquired by Airyscan confocal microscopy. In panel **b**, the scale bar approximately matches the whole cell diameter. In panels **c-e**, the lower scale bar represents the previous cell size, while the upper scale bar represents the current cell size. **f**, Airyscan image of a whole ATTO643-NHS-stained Mega-expanded HeLa cell with a red square highlighting a centriole as an NHS-dense region. **g**, Zoom-in of the centriole marked in (f) single color and Z-color coded projection of the Z-stack. **h,** Mega-ExM Airyscan image of a mitochondrion with its cristae and NPCs as ring-like structures of a primary hippocampal neuron. In addition, a line profile of the cristae signals and a zoom-in into a single NPC ring is shown. All photos were acquired using a 10x 0.45 NA air lens, except for the inset in (b), which was taken with a 40x 1.2 NA water-immersion lens; this is explicitly marked with the respective lens icon in the corner of the image, along with the tile size if a tile acquisition was performed.

**Figure 2.**
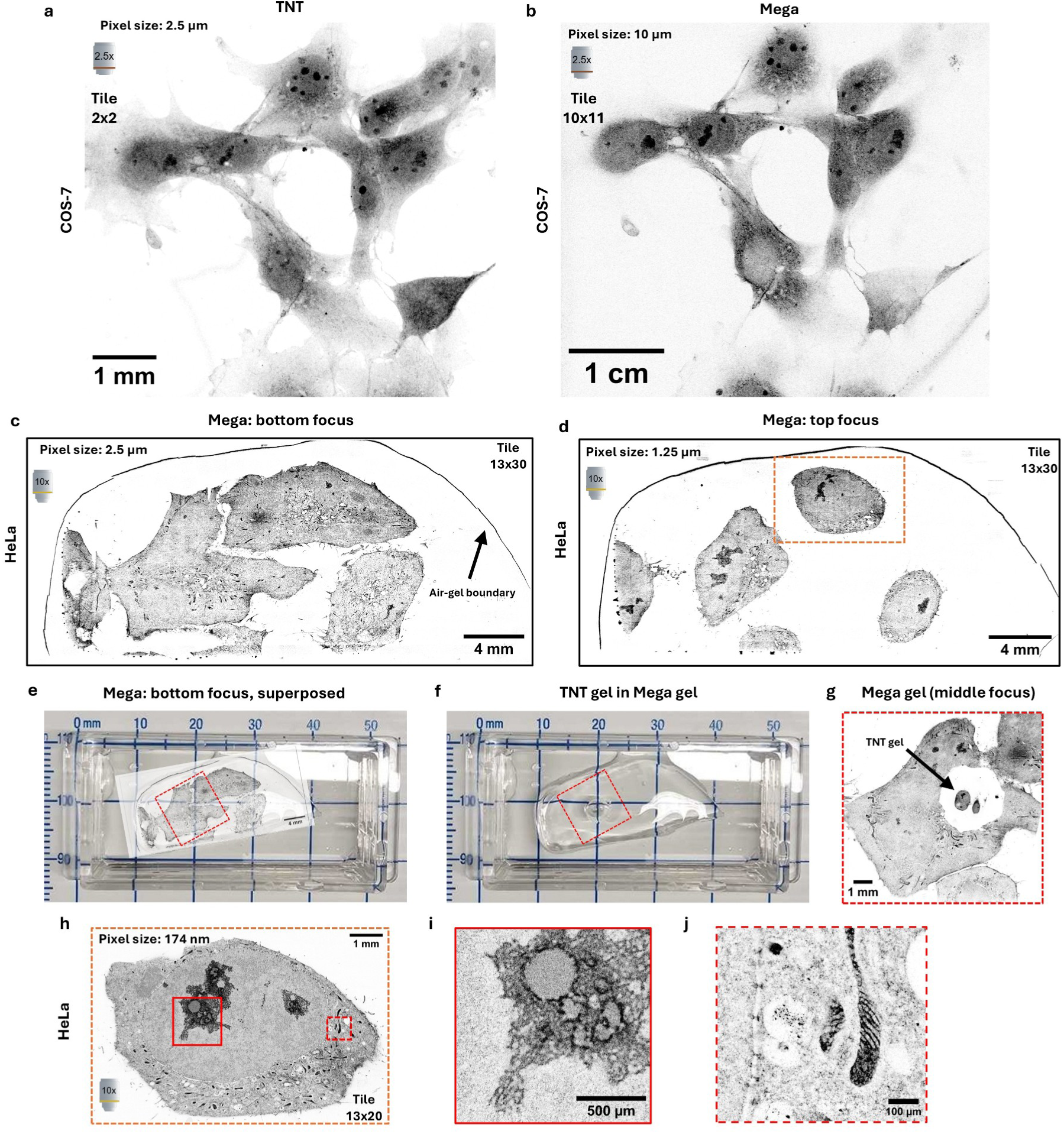
**Size of whole cells in TNT- and Mega-ExM**. **a,b,** Confocal images of the same COS-7 cells labeled with ATTO643-NHS after TNT (∼40x expansion) and Mega (∼200x expansion). **c,d,** Confocal images of Mega-expanded HeLa cells labeled with ATTO643-NHS imaged at different axial planes. **e**, Photo of the Mega-Expanded sample shown in (c,d). **f,g**, Photo and confocal image of a TNT-expanded gel placed in a Mega-expanded gel for size comparison. **h**, Confocal image of a HeLa cell marked by the orange rectangle in (d) imaged at an intermediate focus. **i,j**, Zoom-ins of the red rectangles in (h) showing the nucleolus (i) and mitochondria (j) All acquisition data (lens, pixel size, number of tiles) are shown in the respective images, and all images are shown in inverted grayscale of the ATTO643-NHS staining channel.

We estimated the expansion factor and evaluated distortions by comparing the same cell (or cell region and landmarks) throughout the protocol by Airyscan confocal microscopy, comparing unexpanded cells with expanded cells after one (TREx), two (TNT), and three (Mega) expansion rounds. By simply comparing the size of a HeLa cell before and after each expansion step (Fig. 1b-e), we estimated macroscopic expansion factors of ∼6.6 ± 0.2 (mean ± standard error of the mean, SEM) for TREx, ∼40 ± 1.3 (SEM) for TNT, and ∼228 ± 3.6 (SEM) for Mega-ExM.

For further validation, we plated Raji B-cells on coverslips containing the protein-grid GelMap and applied the Mega-ExM protocol (Supplementary Fig. 6)^30^. Using GelMap we estimated an expansion factor of ∼7.8x in TREx as the 20 µm pattern increased to 155 µm. Using TNT, the distances of the pattern increased to 1 mm, which translates to 50x linear expansion (Supplementary Fig. 6c). We were unable to acquire the GelMap pattern in Mega-ExM gels, as the pattern itself would exhibit a spacing of 5 mm and the signal is strongly diluted, impeding high-quality confocal imaging with low magnification air lenses. B-cells, on the other hand, expanded ∼5.84x ± 0.33 (median ± median absolute deviation, MAD) in TREx, ∼29.0x ± 3.0 (MAD) in TNT, and 249x ± 35 (MAD) in Mega-ExM (Supplementary Fig. 6a-d).

As Mega-expanded cells can have sizes of 0.4-2 cm depending on cell type and spreading on the cover glass, imaging requires the use of low magnification lenses for confocal imaging and several tile images to reconstruct an image of a whole cell (Fig. 1e-f and Fig. 2). Such large expansion factors are suspected to introduce distortions, e.g., through the mounting of the gel on the imaging chamber. However, at first glance, the ultrastructure of mitochondria including cristae, nuclear pore complexes (NPCs), and centrioles appeared to be preserved (Fig. 1f-h). We also applied a fourth round of -NT expansion on already Mega-expanded cells and were able to measure expansion factors of ≥1,500x for “Giga”-expanded B cells (Supplementary Fig. 6e-g). However, the resulting cell dimensions of several centimetres were too large and signal dilution too strong to allow high-end imaging on a conventional confocal microscope (Supplementary Table 3 and Supplementary Fig. 6e).

### Mega-ExM preserves the ultrastructure of macromolecular assemblies in cells

To determine “molecular” expansion factors and compare them between different cellular macromolecular assemblies, we first applied Mega-ExM to the synaptonemal complex (SC). SCs are meiosis-specific multiprotein complexes that are essential for synapsis, recombination and segregation of homologous chromosomes, resulting in the generation of genetically diverse haploid gametes, the prerequisite for sexual reproduction. The SC exhibits an evolutionarily conserved ladder-like organization composed of a central region of ∼114 nm width, plus two lateral elements (∼30-50 nm of width each), to which the chromatin of homologous chromosomes is associated^31,32^. Therefore, the SC has been frequently used as a multiprotein reference structure for super-resolution microscopy and ExM^33–35^. Airyscan confocal microscopy images of immunostained lateral element SYCP3 and dye-NHS labeled SCs clearly confirmed ultrastructure preservation, revealing the ladder-like organization of the SC with a central region width of ∼1.17 ± 0.04 (SEM) µm and ∼6.05 ± 0.31 (SEM) µm after one and two expansion rounds (TNT), respectively (Supplementary Fig. 7). These values correspond to molecular expansion factors of 10.23 ± 0.30 (SEM) and 53.08 ± 2.74 (SEM), respectively. From the measured central region width of 28.96 ± 0.67 (SEM) µm measured from Mega-ExM SC images, we determined a molecular expansion factor of 253.98 ± 6.48 (SEM) for SCs (Supplementary Fig. 7).

As a second macromolecular assembly, we used centrosomes of RPE1 cells and cilia from multiciliated cells from primary hippocampal cultures, which have often been used as rulers in previous ExM experiments^21,23,28^. Centrosomes are membraneless organelles composed of two centrioles, which have a characteristic ninefold microtubule triplet-based symmetry, forming a polarized cylinder ∼400 nm long and ∼200-250 nm wide. The older (mother) centriole has recognizable distal (DA) and subdistal (SDA) appendages, which project radially from the centriole wall and are organized into ninefold-arrays (Fig. 3a)^36–38^. During the G0 phase, the mother centriole serves as a basal body for cilia. 3D Airyscan confocal images of dye-NHS labeled TNT-expanded centrioles allowed the clear identification of their appendages in lateral views (Fig. 3b).

**Figure 3.**
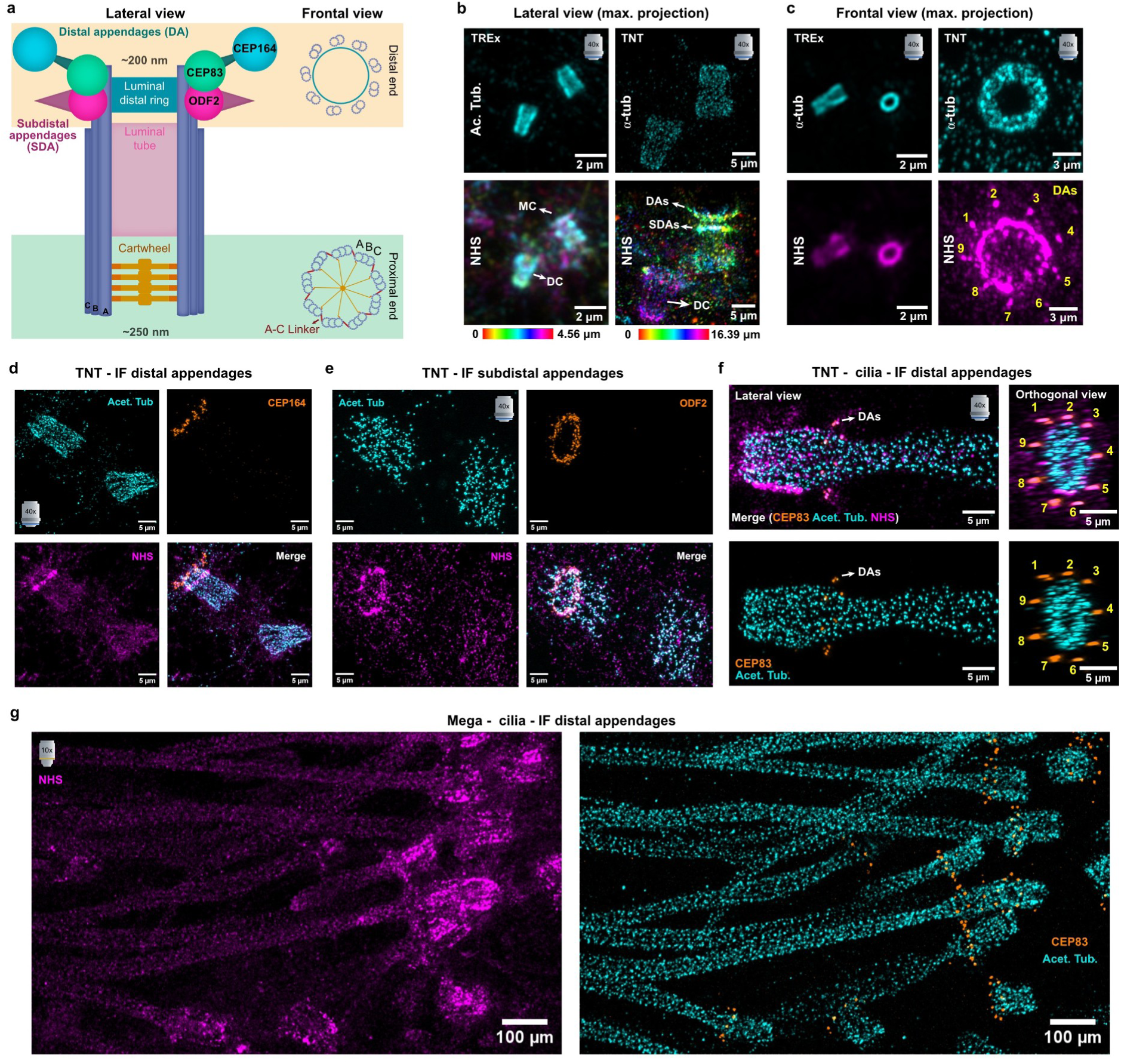
Ultrastructure preservation of TNT- and Mega-ExM in centrosomes and cilia. **a**, depicts the general organization of the mother centriole (MC) in its lateral view, along with schemes of the 9-fold symmetry frontal view where the microtubular doublet (MTD) or triplet (MTT) arrangement can be recognized. The lateral **(b)** and frontal **(c)** views of RPE1 centrosomes (MC and daughter centriole DC) are shown in both TREx and TNT gels stained by NHS and immunostained against α- or acetylated-tubulin. The immunostaining (IF) in TNT of distal (DA) **(d)** or subdistal (SDA) appendages **(e)**, along with acetylated tubulin, are also shown. **f,** An example of cilia in TNT, immunostained against CEP83 (DA) and acetylated tubulin from a multi-ciliated cell in a primary hippocampal culture (putative ependymal multiciliate cell). **g,** Shows a larger field of view of the type of cell shown in **(f)** but in MegaExM. All photos were acquired using a 40X 1.2NA water-immersion lens, except for the MegaExM in **(g)**, where a 10X 0.45NA air lens was used; this is explicitly marked with the respective lens icon in the corner of the images.

In addition, immunostaining of α-tubulin revealed the ultrastructural microtubular organization, including the bundle of nine microtubules with enough resolution to resolve the ninefold triplet/doublet assembly of centrioles in frontal views (Fig. 3c and Fig. 4a-b). Co-immunolabeling with antibodies directed against acetylated-tubulin and CEP164 (DA component) or ODF2 (SDA component) confirmed ultrastructure preservation of centrioles after TNT-expansion (Fig. 3d,e). Centrioles from glial cells in primary hippocampal cultures, and primary cilia of hippocampal neurons also showed good ultrastructure preservation using dye-NHS staining (Fig. 4c-d). Similarly, cilia from multiciliated cells showed no signs of ultrastructural interference after TNT- and Mega-expansion (Fig. 3f,g and Fig. 4g).

**Figure 4.**
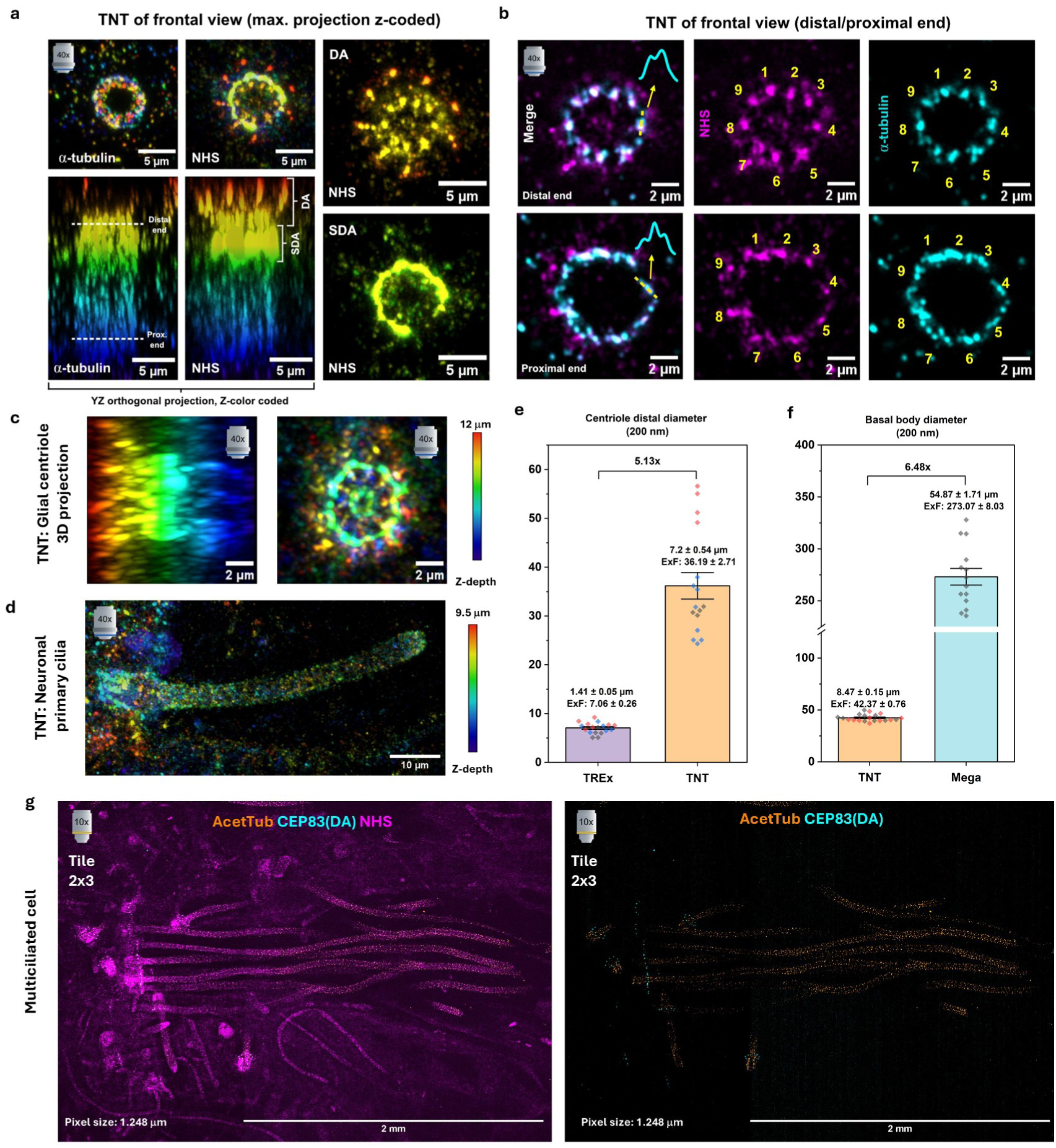
Ultrastructural preservation of centrioles and cilia. a, b, The same centriole presented in the frontal view of Figure 3C for TNT, but with focus on the triplets/doublet assembly of microtubules; with **(a)** showing the projections and orthogonal views for the NHS-staining and a-tubulin channels of the same centriole, highlighting the distal and subdistal appendages visualized by NHS, and **(b)** showing the doublets assembly in the distal end of the centriole and the triples arrangement of the proximal end. **c, d**, Glial centriole and a neuronal primary cilium respectively from ATTO643-NHS-stained primary hippocampal cultures, in TNT expansion stage. **e, f,** The estimation of the expansion factors of TREx, TNT and Mega by centriole **(e)** and basal body **(f)** distal-end diameter (200 nm) is shown. **g**, depicts a tile image of Mega from multiciliated cells from the primary hippocampal cultures, identified by the immunostaining against acetylated tubulin and the distal appendages protein CEP83, depicting cilia that extends beyond the 2 mm in length. For **(e)**, values were measured from three independent experiments (N=3, dots color-coded), with a mean diameter of 1.41 ± 0.22 µm for TREx (18 centrioles: n1=5, n2=7, n3=6) and 7.24 ± 2.17 µm for TNT (16 centrioles: n1=4, n2=4, n3=8). For **(f)**, diameters were measured for two independent experiments for TNT (N=2, dots color-coded) and one from Mega (N=1), with mean diameters of 8.47 ± 0.15 µm for TNT (20 cilia: n1=13, n2=7) and 54.61 ± 1.61 µm for Mega (n1=14). Data presented as Mean ± SEM in µm.

From centrosomes, we used the mother centriole’s diameter on its distal end to estimate a mean molecular expansion factor of 7.1 ± 0.3 (SEM) for TREx and 36.2 ± 2.7 (SEM) for TNT expansion (Fig. 4e). For Mega expansion, an expansion factor was measured for multiciliated cells, with basal body diameters of 54.6 ± 1.6 µm (SEM) corresponding to an expansion factor of 273.1 ± 8.0 (SEM, Fig. 4f). For comparison, in TNT-expanded centrioles the diameter was measured to 8.5 ± 0.2 µm (SEM), which translates to an expansion factor of 42.4 ± 0.8 (SEM). Furthermore, multiciliated cilia in the brain rank among the longest cilia in mammals, with a typical length of approximately 8 µm^39^. In Mega-ExM they exhibit lengths of >2 mm (Fig. 4g), which further confirms an expansion factor that surpasses 200x.

### Mega-ExM resolves mitochondrial ultrastructure

To further demonstrate the performance and validate the expansion factor achievable by Mega-ExM, we investigated ultrastructural key features of mitochondria. Mitochondria are double-membrane organelles that serve as the central hubs of cellular energy metabolism in nearly all eukaryotes^40^. Rather than forming a uniform membrane sheet, the inner membrane is extensively folded into highly curved, dynamic invaginations termed cristae, which dramatically expand membrane surface area and create distinct biochemical microcompartments within the organelle^41^. As revealed by cryo-electron tomography, the crista-to-crista spacing (i.e., the border-to-border distance) in mammalian cells is typically 30 –100 nm, whereas the membrane-to-membrane spacing within lamellar cristae (i.e., the crista width) is much smaller (20–40 nm) and relatively well preserved. (Fig. 5a)^42,43^. Murine neuronal hippocampal mitochondria exhibit very tight crista spacing, with a median minimum distance of ∼28 nm, and a cristae width of ∼36 nm, positioning them at the low end of reported crista-to-crista distances (Fig. 5a)^44^. Therefore, we selected primary hippocampal cultures to assess the ability of Mega-ExM to resolve cristae in mitochondria.

**Figure 5.**
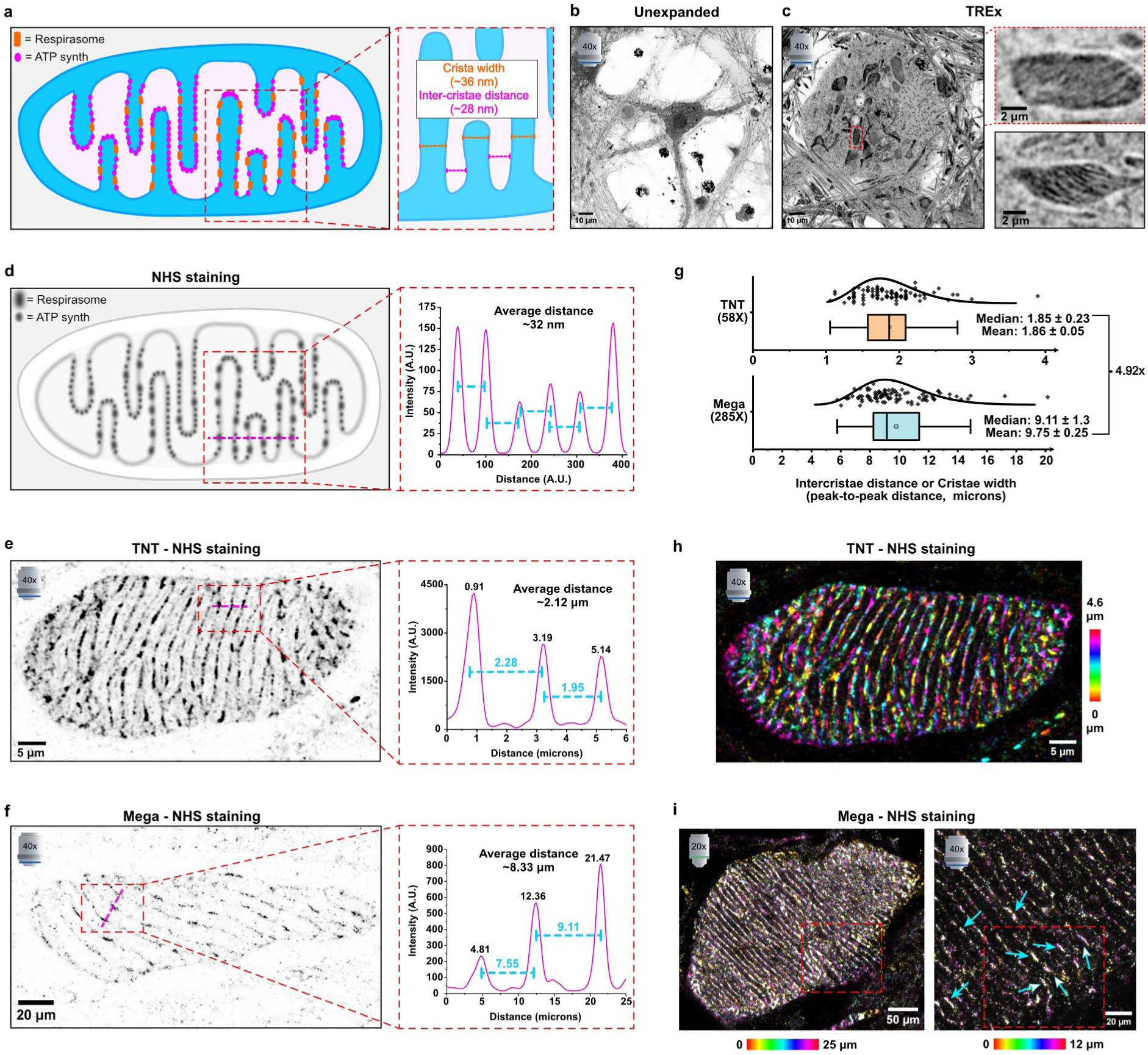
Cristae-to-cristae spacing and cristae width in neurons as internal molecular ruler for Mega-ExM expansion factor estimation. a, Model on how cristae width and inter-cristae distance are measured typically in EM. **b,c,** Airyscan images of ATTO643-NHS labeled unexpanded and TREx-expanded mouse primary hippocampal neurons. The unexpanded image shows only one neuronal soma in the field of view. In TREx-expanded samples inter-cristae spacing can be occasionally resolved. **d,** Model explaining how cristae-to-cristae spacing was analyzed in expanded dye-NHS-stained mitochondria. The average inter-cristae distance was determined by measuring the distance between the signal maxima of different cristae, which should result in an average distance of ∼32 nm for mitochondria in primary hippocampal neurons. **e,f**, Airyscan images of TNT- and Mega-expanded mitochondria of primary hippocampal neurons and corresponding inter-cristae analysis. **g**, Distribution of 92 and 93 distances measured in TNT- and Mega-ExM for 10 and 11 different mitochondria, respectively, each across three independent experiments. Inter-cristae distances are given as Mean +/- SEM and Median +/- MAD. Two independent experiments for Mega- and three for TNT-ExM (different primary culture and ExM experiment) were performed. **h,** Airyscan image of a z-color-coded projection of a TNT-ExM mitochondrion. **i,** Airyscan images of z-color-coded projections of a Mega-ExM mitochondrion imaged with a 20x 0.85 N.A. air objective (left) and a 40x 1.2 N.A. water immersion objective (right) where arrows indicate individual cristae protein density signals (arrows). The red rectangle of the right panel is the field of view later shown in Figure 7.

Structural details of cristae are too small to be resolvable by standard confocal microscopy and TREx in combination with Airyscan confocal microscopy, which provides a spatial resolution of ∼30 nm on ∼8x expanded samples (Fig. 5b,c)^45^. On the contrary, Airyscan microscopy of dye-NHS labeled TNT-expanded mitochondria should allow determining cristae spacing and width in hippocampal neurons (Fig. 5a,d). Peak-to-peak distances between cristae ranged from 1.6-2.1 µm in TNT-expanded mitochondria and from 7.6-10.7 µm in Mega-expanded mitochondria (Fig. 5e-g). Of note, although mitochondria from hippocampal neurons require at least TNT to fully resolve the separation between cristae, a single TREx is sufficient for COS-7 and HeLa cells, where the inter-cristae distance is reported to be in the 70-100 nm range (Supplementary Fig. 8)^42,43^.

Assuming an average inter-cristae distance of 32 nm (Fig. 5a,d) we determined expansion factors of 57.8 ± 7.2x (MAD) for TNT-ExM and 284.7 ± 40.6x (MAD) for Mega-ExM (Fig. 5g). Because the EM comparison metrics gives minimum inter-cristae distance and minimum cristae width, whereas our line-profile measurements capture protein-density peak-to-peak spacing rather than direct membrane segmentation, our values likely overestimate the true expansion factors but can be used as an approximation. For example, using the SC-derived expansion factors of ∼50x (TNT) and ∼250x (Mega) as reference (Supplementary Fig. 7d), we determined inter-cristae distances of ∼37 nm and ∼36 nm, respectively, which remain within the expected range for mitochondria of primary hippocampal neurons^44^.

Overall, our results show that each additional expansion round of TREx-expanded samples increases the expansion factor of macromolecular assemblies and organelles by at least 5-fold (Fig. 4c,d, Fig. 5g, and Supplementary Fig. 7d). This is consistent with a shrinkage of ∼70-80% observed for NT re-embedding, which reduces the final practically achieved expansion factor of the TREx gel in the second and third expansion round (Supplementary Table 2).

When inspecting color-coded z-projections of ATTO643-NHS-labeled expanded mitochondria in more detail, it becomes apparent that Mega-ExM can resolve cristae curvatures, mitochondrial ultrastructure, morphology and interconnections using conventional air lenses (Fig. 5h,i and Fig. 6). Hence, Mega-ExM could also be applied to study changes in morphology of mitochondrial cristae that have been linked to different diseases including cancer, diabetes, aging, and neurodegenerative diseases^46,47^. Furthermore, Mega-ExM reveals protein-dense regions located on flat membrane regions of cristae, possibly corresponding to respiratory chain supercomplexes (respirasomes) (Fig. 5i)^48^.

**Figure 6.**
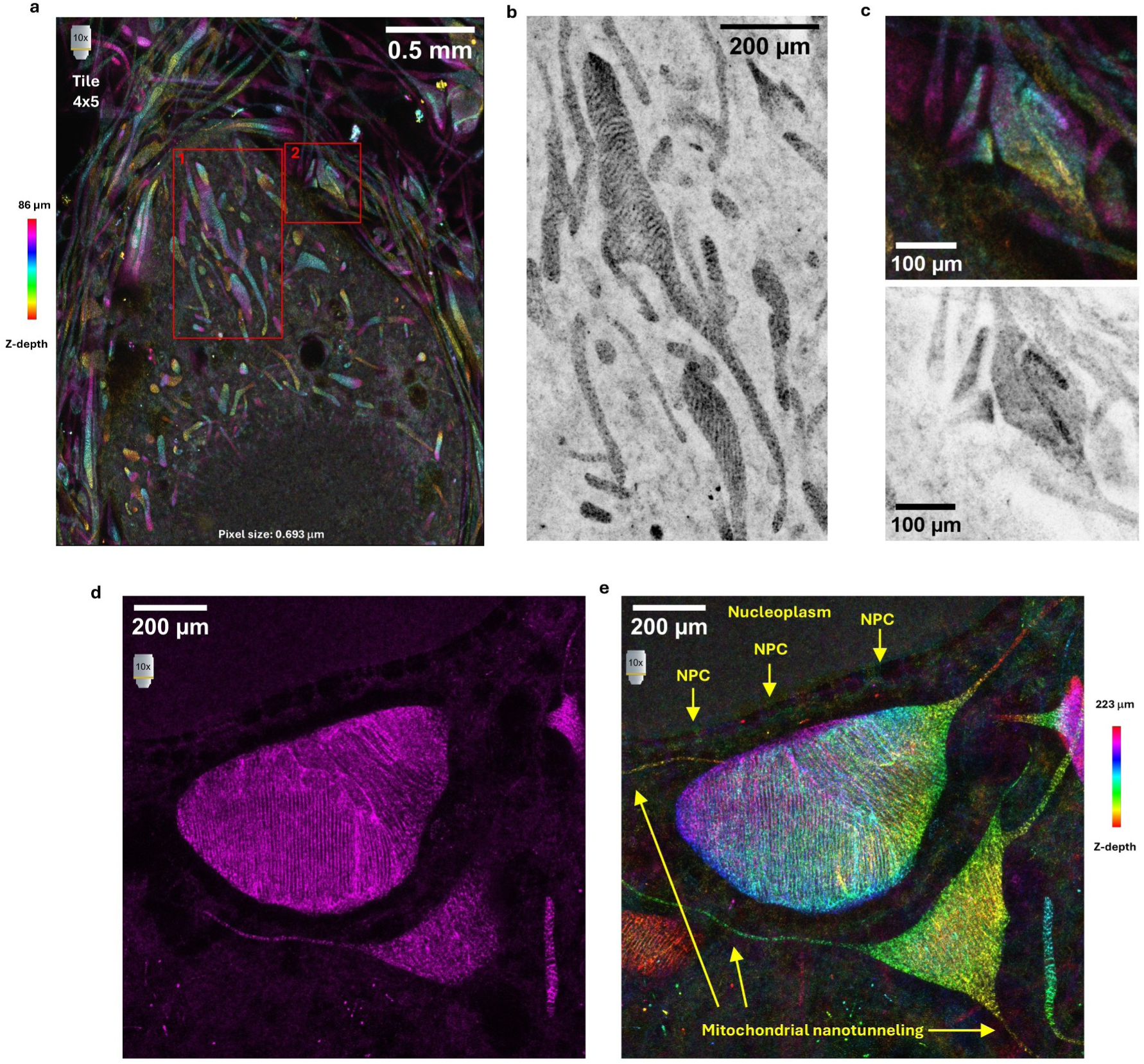
**Mega-ExM unlocks cellular and mitochondrial ultrastructure using conventional air objectives. a-c**, Airyscan images of an individual hippocampal neuron soma from a tile z-stack (a), showing the nuclei at the bottom with NPCs visualized as dense dots in the nucleoplasm/cytoplasm border, and a primary dendrite extending in the top part of the image covered with dendrites/axons (see also Supplementary Video 3). Zoom-ins into the red rectangles highlight mitochondria (b) and synapses (c), respectively using a conventional 10x air objective. An under-sampled tile (693 nm pixel size compared to the 174 nm for Airyscan and 348 for normal confocal) is sufficient to resolve the 30 nm inter-cristae distances of hippocampal neurons (b) and single synapses with its synaptic cleft of ∼30 nm (c). **d**, Airyscan image of the soma of a hippocampal neuron with focus on the mitochondrial network near the nuclei (upper part of the image). **e**, Color-coded z-projection of the same sample annotated to highlight NPCs in the nucleoplasm/cytoplasm border in side view, along with the extensions of mitochondria forming nanotunnels for inter-mitochondria communication.

Cristae are not merely architectural adaptations to increase the surface area; they play an active role in organizing and regulating the respiratory machinery. The respiratory complex, a supercomplex comprising several membrane-embedded molecular machines, generates proton motive force, which in turn drives ATP synthase^49,50^. Although high-resolution structures of isolated respiratory complexes and respirasomes have provided important mechanistic insights, their supramolecular arrangement within intact cristae remains incompletely understood^51,52^. Respirasomes are large mitochondrial supercomplexes composed primarily of Complex I, a dimer of Complex III, and one or more copies of Complex IV. Their total molecular mass is typically in the range of ∼1.5–2.0 MDa, making them substantially larger than individual respiratory complexes, and structural studies indicate dimensions on the order of ∼20-30 nm^48,51,52^.

Recently, cryo-electron tomography resolved native structures of respiratory chain complexes and revealed how respiratory complexes I, III, and IV assemble into a respirasome supercomplex, which is restricted to flat membrane regions apart from rows of ATP synthase at the curved tips of cristae^53^. Since Mega-ExM provides images of mitochondria that reveal the organization of cristae, we zoomed into these cristae structures using a water immersion lens instead of air lenses (Fig. 5i) to verify if the achieved expansion factor and NHS-labeling density allows us to visualize respirasomes in their native cristae context by Airyscan confocal microscopy. Therefore, we identified protein-dense regions on cristae of TNT- and Mega-expanded mitochondria in primary hippocampal neurons that exhibit a length of ∼30 nm, matching the size of individual respirasomes (Fig. 7a,b)^48,51–53^.

**Figure 7.**
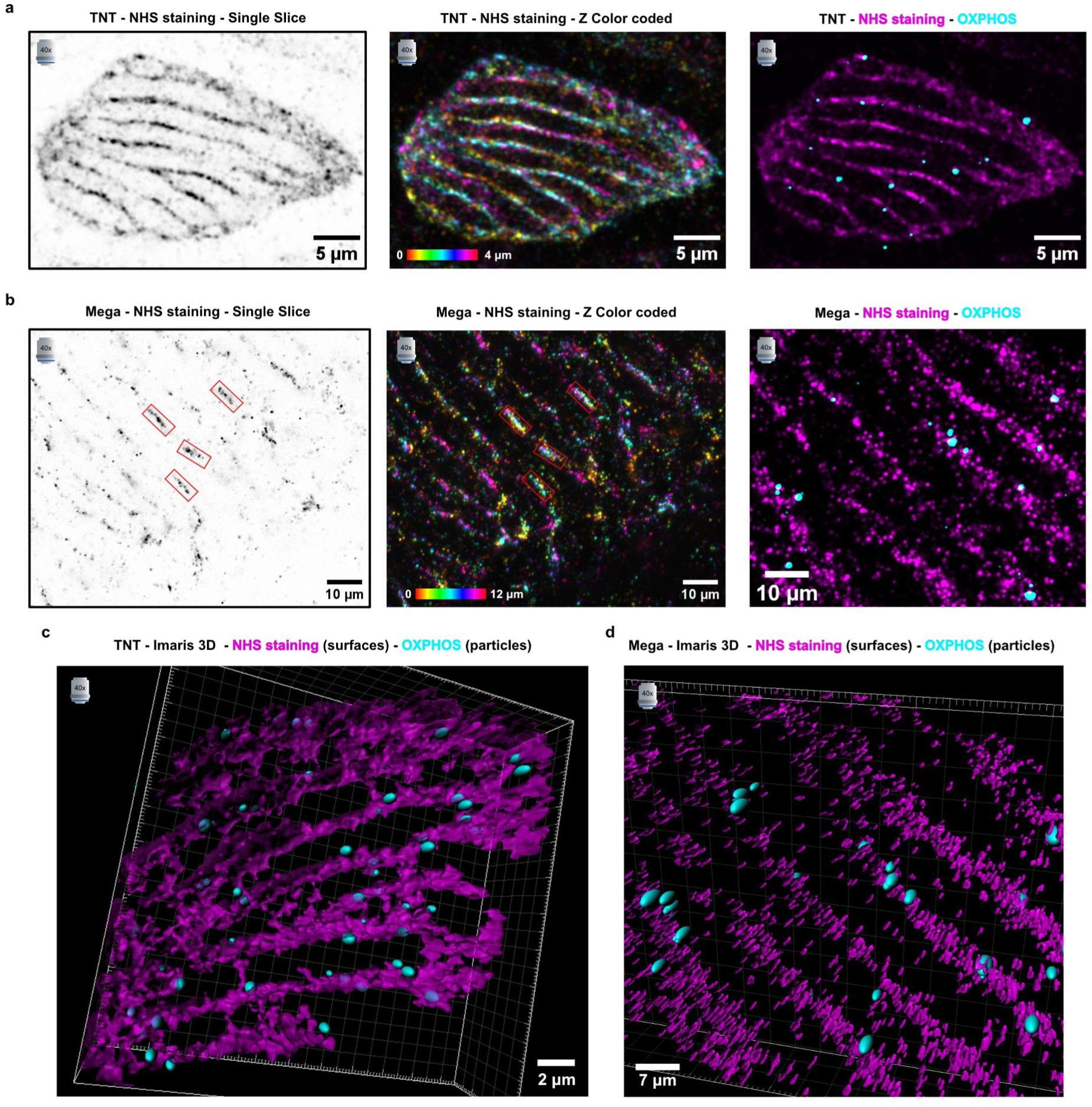
OXPHOS and respirasome localization in mitochondrial cristae by Mega-ExM. a, Airyscan image of a TNT-expanded mitochondrion of a primary hippocampal neuron stained with ATTO643-NHS followed by immunolabeling for OXPHOS components using a cocktail of five monoclonal antibodies (cyan). **b,** Airyscan image of the red rectangle depicted in the Mega-ExM mitochondrion shown in Fig. 5i. In both the inverted grayscale (left) and the z-color-coded (middle) panels, small red rectangles are surrounding putative respirasome; the size of the rectangle was chosen to be 37 nm x 15 nm considering an expansion factor of 250x, corresponding to the size of a single respirasome viewed from the side^52^. An example of subsequent immunostaining against OXPHOS is shown on the right. **c,d,** Rendered 3D image of the mitochondria shown in (a) and, with a magenta surface representation for dye-NHS staining, and cyan spheres for the localization of the immunolabeled OXPHOS signal.

And, in fact, protein-dense structures identified in Mega-ExM on flat membrane regions of cristae correspond well in size to the respirasome structures visualized by cryo-electron tomography (Fig. 7b)^53^ considering 250x expansion. Simultaneous immunolabeling of the whole OXPHOS components (Complex I-V) from a commercial cocktail of monoclonal antibodies against individual complex proteins confirmed that the majority of clustered and dense NHS-signals on flat membrane areas of cristae contain OXPHOS components (Fig. 7b-c).

### Mega-ExM of nuclear pore complexes challenges cryo-electron tomography

To demonstrate that Mega-ExM can approach the resolution achieved by cryo-electron tomography, we labeled Hela cells, COS-7 cells, and neurons with ATTO643-NHS and focused on visualizing nuclear pore complexes (NPCs) embedded in the nuclear envelope. The NPC is one of the largest supramolecular assemblies in eukaryotic cells that regulates bidirectional traffic of macromolecules across the nuclear envelope. The NPC exhibits an eight- and twofold rotational symmetry across the nuclear envelope plane and along the nucleocytoplasmic axis, respectively^54,55^. To optimize ultrastructure preservation of NPCs we first investigated different combinations of acrylamide (AA) and formaldehyde (FA) for crosslinking and anchoring, respectively, and denaturation conditions. Here, 2% AA and 1.4% FA combined with 85°C denaturation at pH 6.8 provided the best results (Supplementary Fig. 9). To improve the image quality of TNT- and Mega-expanded samples, we sliced the expanded hydrogels into smaller segments to enable Airyscan confocal imaging with a 40x NA 1.2 water-immersion objective. Images of the nuclear membrane of TREx-expanded Hela cells and neurons showed NPCs as densely labeled signals (Fig. 8a and Fig. 9a). In contrast, Airyscan images of TNT-ExM processed Hela cells, neurons and COS-7 cells, disclosed the ring-like structure of NPCs and their eightfold symmetry (Fig. 8b-c, Fig. 9b-c and Supplementary Fig. 10a).

**Figure 8.**
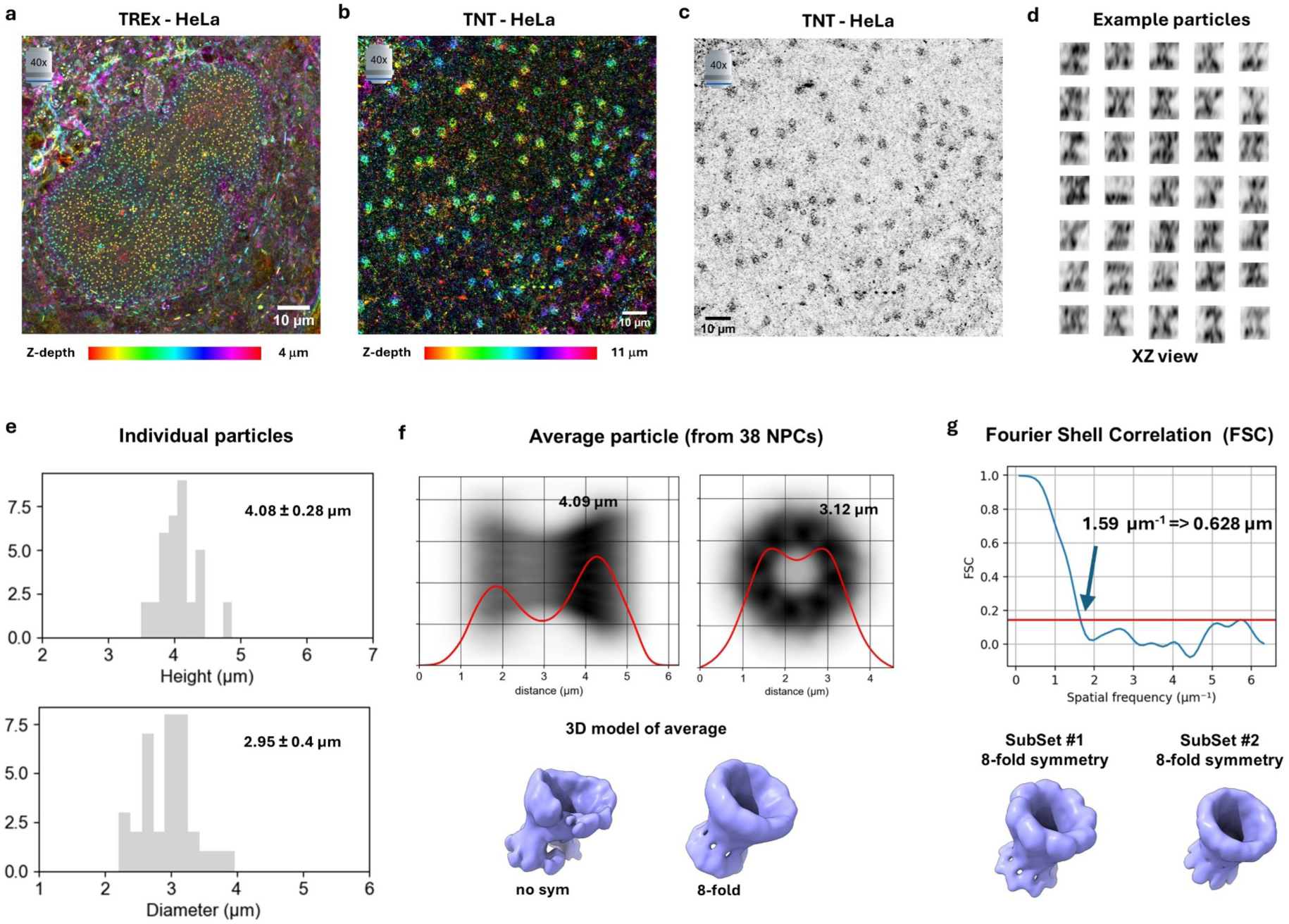
NPC reconstruction by particle averaging of ATTO643-NHS labeled TNT-expanded HeLa cells. **a**, 3D-Airyscan image of a TREx gel of a HeLa cell, with focusing on the bottom of the nuclei, shown as z-color-coded projection, where NPCs can be visualized as dense dots (see also Supplementary Figure 9b). **b**, 3D-Airyscan Image of a TNT gel of (a), where NPCs are now resolved as circular ring-like structures. **c**, Maximum intensity projection of (b) shown as inverted scale for improved visualization of NPC rings. **d**, Examples of isolated particles (individual NPCs) in their side (*xz*)-view, where at least two rings (cytoplasmic and nuclear) can be recognized. **e**, Histograms for the diameter of rings and distances between rings (height) detected for individual NPCs. **f**, Reconstruction of an average NPC using Fourier correlation alignment of N=38 particles for a single field of view of one nucleus of a HeLa cell, with 8-fold symmetry imposed, shown as side (zx, left) and frontal (xy, right) view sum projection with intensity profiles (red) superimposed. The lower panel shows 3D reconstructions of averaged particles with and without 8-fold symmetry. **g**, Estimation of the practical resolution achieved in reconstructed average NPC structures by Fourier Shell Correlation (FSC) of two datasets of NPCs acquired from the same gel and same HeLa cell.

**Figure 9.**
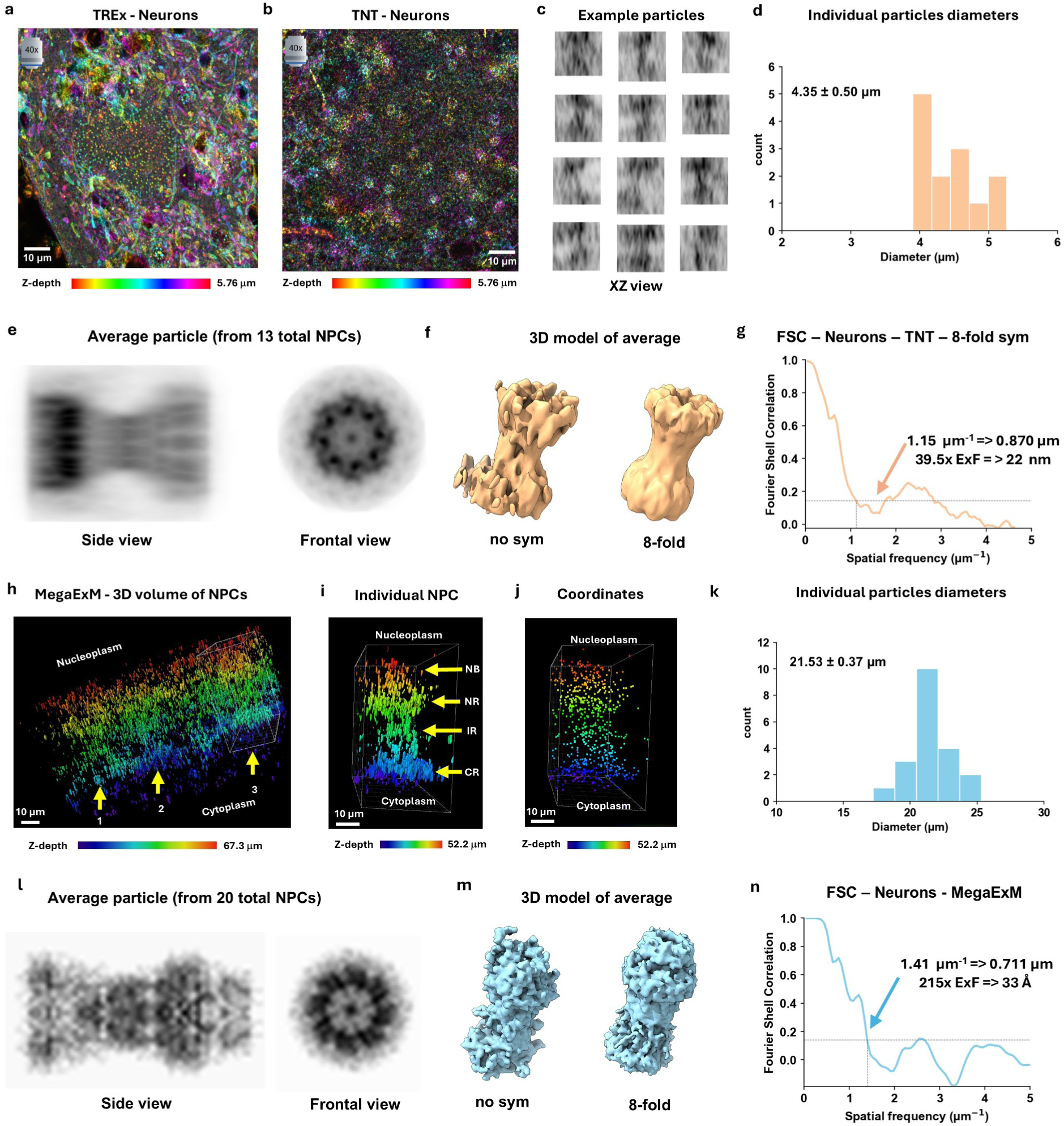
**NPC reconstruction by particle averaging of ATTO643-NHS labeled TNT- and Mega-expanded hippocampal neurons**. **a**, 3D-Airyscan image of a TREx gel of a primary hippocampal neuron focusing on the bottom of the nuclei, shown as z-color-coded projection, where NPCs can be visualized as dense dots (see also Supplementary Figure 9b). **b**, 3D-Airyscan image of a TNT-expanded hippocampal neuron, where NPCs are resolved as circular ring-like structures. **c**, Examples of isolated particles (individual NPCs) in their *xz* view, showing the cytoplasmic and nuclear ring. **d**, Histogram of NPC diameter rings determined from individual TNT-expanded NPC particles. **e**, Reconstruction of an average NPC using Fourier correlation alignment of N=13 particles for a single field of view of one nucleus of a hippocampal neuron, with 8-fold symmetry imposed, shown as side view (*zx*, left) and frontal view (*xy*, right) projection. **f**, 3D reconstruction of averaged NPC particles with and without 8-fold symmetry imposed. **g,** Estimation of the practical resolution achieved in reconstructed average NPC structures by Fourier Shell Correlation (FSC) of two datasets of NPCs acquired from the same gel and same neuron. **h,** 3D Airyscan images of five Mega-expanded NPCs in hippocampal neurons shown in side view, where three NPCs are easily visualized (identification is facilitated by z-color-codes). **i**, Single particle (NPC) number #3 isolated from (**h**) showing individual fluorescence signals (PSFs). Nevertheless, structural features of the NPC including nuclear- (NR), cytoplasmic- (CR), and inner rings (IR), in addition to the nuclear basket (NB) are readily visible. **j**, 3D-positions of the fluorescence signals of NPC were reduced to coordinates for particle-averaging, as shown here for particle #3. **k**, Histogram of NPC diameter rings determined from individual Mega-expanded NPC particles. **l**, Reconstruction of an average NPC using Fourier correlation alignment of N=20 Mega-expanded particles using the localization coordinates of their individual signals, with 8-fold symmetry imposed, shown as side view (*zx*, left) and frontal view (*xy*, right) projection. **m,** 3D reconstruction of averaged NPC particles with and without 8-fold symmetry imposed. **n,** Estimation of the practical resolution achieved in reconstructed average NPC structures by FSC of two datasets of Mega-expanded NPCs from neurons (10 each). Values for TNT are from the dataset identified as Neuron-1-1-1 (first column of Supplementary Table 4) and given as Mean +/- SD, while Mean +/- SEM was used for the diameters in (k).

Cryo-electron tomography (cryo-ET) has emerged as a key approach for elucidating the structure of large multiprotein assemblies such as the nuclear pore complex (NPC) within their native cellular context. In tomograms, NPCs appear as ring-shaped densities spanning the double membrane of the nuclear envelope. Because individual tomograms are inherently noisy and limited by the missing-wedge artifact, sub-tomogram averaging is used, i.e., hundreds of NPC particles are extracted, aligned according to their conserved eightfold symmetry, and averaged to enhance the signal-to-noise ratio^56,57^. Applying cryo-ET to intact human fibroblasts and HeLa cells allowed the reconstruction of mammalian NPCs at 35-66 Ångström resolution^57–59^. To compare our results of TNT-expanded NPCs with those of cryo-ET, we selected particles, determined their individual diameters, and applied particle averaging with and without imposing an 8-fold symmetry to reconstruct NPC images with improved signal-to-noise ratio (Fig. 8d-f, Fig. 9c-f, and Supplementary Fig. 10b-f).

From the average reconstructed images, we determined NPC ring diameters and distances between the cytoplasmic and nuclear rings of 4.2 µm and 7.3 µm in hippocampal neurons, 2.5 µm and 4.4 µm in Hela cells, and 2.7 µm and 4.9 µm in COS-7 cells (Fig. 9e,f, Fig. 8e,f, Supplementary Fig. 10d,e, and Supplementary Table 4). These values translate into expansion factors of ∼37.8x for hippocampal neurons, ∼22.6x for Hela, and ∼24.3x for COS-7 cells, assuming an NPC diameter of ∼110 nm^56,57^, as a conservative approach for labelling with NHS-dyes. Consistent with previous reports, denaturation-based expansion yielded larger expansion factors along the nuclear transport axis, particularly for NPCs in neurons^60,61^.

Using Fourier Shell Correlation (FSC) analysis^62,63^ we estimated an average conservative resolution of ∼830 nm for the NPC reconstructions from TNT-expanded hippocampal neurons (Fig. 9g, and Supplementary Table 4), corresponding to a potential structural resolution of ∼22 nm in unexpanded dimensions. Similarly, we estimated a structural resolution obtained from particle averaging of TNT-expanded NPCs of ∼23 nm and ∼28 nm in Hela and COS-7 cells, respectively, after correcting for the expansion factor (Fig. 8g, Supplementary Fig. 10g and Supplementary Table 4).

Motivated by these results, we investigated Mega-ExM Airyscan images of NPCs in hippocampal neurons, as they exhibited the largest expansion factors following TNT expansion. Here, NPCs appear as protein assemblies with a diameter of ∼20 µm corresponding to an expansion factor of ∼200x (Fig. 9h). At this level of expansion, individual point spread functions (PSFs), likely corresponding to individual fluorescently labeled NPC proteins, can be resolved in 3D Airyscan images. Nevertheless, these images reveal the overall architecture of the NPC, including the nuclear, cytoplasmic, and inner rings, as well as the nuclear basket (Fig. 9i). Therefore, we used the localization coordinates of individual fluorescence signals for particle averaging as previously described for expansion microscopy-based analysis of individual protein structures (Fig. 9j)^5^. Using the measured NPC mean diameter of 21.528 ± 0.374 µm (SEM) we estimated an expansion factor of ∼215x for Mega-ExM of NPCs (Fig. 9k). Using only 20 NPC particles for averaging allowed us to reconstruct a 3D model of the NPC with a putative structural resolution of ∼33 Ångström (Fig. 9l-n). This resolution challenges the structural resolution achieved by cryo-ET for human NPCs from HeLa^57^ and human fibroblast cells^59^.

Interestingly, Mega-ExM resolved structural features of the NPC nuclear basket that are less apparent in cryo-ET reconstructions (Fig. 9l-m). This apparent discrepancy reflects the fundamentally different contrast mechanisms of the two techniques. Mega-ExM visualizes the distribution of surface-exposed lysine residues through NHS-dye labeling, whereas cryo-ET derives contrast from electron density. Consequently, although cryo-ET generally provides higher structural resolution, flexible and weakly electron-dense protein assemblies remain challenging to resolve^58,64^. By contrast, lysine-rich protein structures that are efficiently labeled with NHS-functionalized dyes can be visualized with remarkable detail by Mega-ExM, allowing the method to approach the structural resolution achieved by cryo-ET for these specific assemblies.

### Mega-ExM and labeling imperfections

Our data show that the labeling density of aliphatic amino groups of Mega-expanded NPCs and mitochondrial proteins with the dye ATTO643-NHS is high enough to enable fluorescence imaging on an Airyscan microscope with a structural resolution of several tens of Ångström in cells. However, other cellular structures are not labeled with ATTO643-NHS at a density required to achieve similar resolutions, and some cellular structures including microtubule and actin filaments are not visualized by Mega-ExM. Basically, there are two mechanisms that can prevent the visualization of proteins by Mega-ExM: either the proteins are not efficiently anchored into the hydrogel and/or they are not efficiently labeled by NHS-functionalized dyes. In addition, visualization is affected by the anchoring denaturation conditions. Microtubules, for example, are prone to depolymerization due to inefficient fixation and poor anchoring of proteins into the hydrogel, which causes visualization problems as encountered in early EM work^65^. Similarly, actin filaments are only inefficiently anchored into the hydrogel and thus difficult to visualize by ExM^66^.

Furthermore, the physicochemical properties of the dye itself may influence the labeling efficiency of specific cellular structures. For example, hydrophobic cationic dyes can diffuse across cellular membranes and preferentially accumulate in mitochondria because of the large negative membrane potential across the inner mitochondrial membrane^67^. Whether this preferential accumulation is preserved in expanded hydrogels, however, and to what extent it influences staining patterns, remains unclear. To investigate the impact of the hydrophobicity of the dye structure on the labeling selectivity, we performed expansion experiments using 13 different NHS-functionalized dyes^68^. We evaluated at the TREx stage the most recognizable compartments/organelles with our protocol (nucleolus, NPCs, mitochondria, centrosomes, chromosomes and nucleoplasm) and combined different NHS-functionalized dyes to find out whether they compete for labeling cellular structures (Supplementary Figs. 11-14 and Supplementary Table 5). In summary, more hydrophilic dyes such as CF dyes, Alexa Fluor 488, ATTO488 and ATTO643, tend to produce intense labeling of nucleoli, nucleoplasm and centrosomes but comparatively weak or moderate mitochondrial staining. In contrast, more lipophilic and hydrophobic cationic dyes such as Cy3, ATTO550, ATTO633, ATTO647N, and ATTO665 show pronounced labeling of mitochondria and nuclear pore complexes while still efficiently labeling nucleoli and centrosomes. Notably, even dyes with similar manufacturer-reported hydrophilicity (for example, Alexa Fluor 488, CF555 and CF568, or the different far-red ATTO dyes) exhibited distinct labeling patterns in side-by-side competition experiments, indicating that subtle differences in dye structure can modulate the labeling efficiency of cellular organelles (Supplementary Figs. 11-14 and Supplementary Table 6). These findings suggest that the current view of covalent labeling in cells should be expanded to account not only for the reactivity of the functional group but also for the physicochemical properties of the fluorophore, including its hydrophobicity and charge. This raises the possibility of designing NHS-functionalized dyes that preferentially label distinct intracellular structures.

## Discussion

Expansion microscopy (ExM) has transformed fluorescence imaging by enabling nanoscale visualization of cells and tissues using conventional diffraction-limited microscopes. By physically enlarging specimens before imaging, ExM provides straightforward access to super-resolution information without requiring specialized instrumentation or extensive technical expertise. Initial concerns regarding the fidelity of the expansion process, particularly isotropy and ultrastructural preservation, have largely been resolved through optimized protocols and rigorous validation strategies^12–15,17–22^. Nevertheless, translating the theoretical resolving power of ExM into molecular-scale structural imaging has remained challenging because substantially higher expansion factors must be combined with efficient fluorescence labeling and sensitive fluorescence detection^5,6,24,25^. Recently, thousandfold expansion microscopy (1000ExM) demonstrated that resolving adjacent lysine residues in isolated proteins and peptides is possible by combining extreme expansion with single-molecule-sensitive confocal microscopy. However, this approach relies on proteinase K digestion of anchored proteins before expansion, making it unsuitable for preserving the ultrastructure of macromolecular assemblies in intact cells at high expansion factors^27^.

Mega-ExM does not have this limitation as it uses heat-denaturation homogenization instead of enzymatic digestion. By using three TREx expansion rounds interspersed with two stabilizing, cleavable neutral hydrogels, Mega-ExM preserves protein integrity while achieving ∼250-fold expansion of macromolecular complexes and organelles in their native cellular context. Although expansion factors vary slightly between structures, ultrastructural organization is faithfully preserved, enabling fluorescence imaging of entire cells with unprecedented structural resolution using conventional confocal microscopes.

Structural characterization of macromolecular complexes has traditionally relied on *ex-situ* approaches using purified samples, most notably cryo-electron microscopy. However, many protein assemblies require their native cellular environment for correct assembly, organization, and function. For example, nuclear pore complexes (NPCs) form only within the context of the fused inner and outer nuclear membranes^56^. Cryo-electron tomography (cryo-ET) has therefore become the method of choice for visualizing macromolecular architecture *in-situ*^58,64^. Remarkably, Mega-ExM images of NPCs closely resemble corresponding cryo-ET reconstructions. For NPCs in neurons, particle averaging yielded a structural resolution for the reconstructions of ∼35 Å, demonstrating that, under favourable conditions, Mega-ExM can approach the structural resolution achieved by cryo-ET. Importantly, the two techniques derive image contrast from fundamentally different physical properties. Whereas cryo-ET visualizes electron density, Mega-ExM reports the spatial distribution of accessible lysine residues through NHS-dye labeling. Consequently, the structural resolution attainable by Mega-ExM depends primarily on labeling density and the accessibility of lysine residues rather than on electron density.

Although Mega-ExM samples can be expanded beyond 1,000-fold using our established protocol (“Giga-ExM”), the resulting cell dimensions (2-5 cm), together with fluorophore dilution and the accompanying reduction in signal-to-noise ratio, currently limit imaging at higher structural resolutions using conventional confocal microscopes. Consequently, expansion factors and labeling density must be carefully balanced to achieve optimal structural resolution. The optimal expansion factor depends on the complexity of the protein assembly and, critically, on the density and accessibility of reactive lysine residues, which vary substantially among cellular structures.

Further improvements in structural resolution will therefore depend primarily on advances in labeling chemistry rather than expansion chemistry. Because lysine residues constitute only ∼6-8% of amino acids, NHS labeling is approaching its theoretical labeling-density limit. More reactive functional groups capable of targeting additional biomolecular classes including proteins, lipids, carbohydrates, and nucleic acids could substantially increase labeling density and thereby exploit higher expansion factors. Likewise, tailoring the physicochemical properties of fluorophores or coupling them to affinity motifs may enable selective labeling of specific intracellular structures. Combined with probes targeting different amino acids, such advances could ultimately enable multicolor Mega-ExM for spatially resolved mapping of protein composition directly within intact cells.

In summary, Mega-ExM establishes fluorescence microscopy as a structural imaging modality capable of visualizing macromolecular architecture in intact cells. By combining molecular specificity with structural resolution approaching that achieved by cryo-ET for selected protein assemblies, Mega-ExM complements electron microscopy while remaining compatible with conventional fluorescence microscopes, millimeter-scale specimens, and established image-analysis workflows. Rather than representing simply another expansion microscopy protocol with a larger expansion factor, Mega-ExM provides a broadly accessible platform for structural cell biology. As labeling chemistry and probe design continue to advance, Mega-ExM has the potential to bridge molecular specificity, ultrastructure, and spatial biology within the same specimen, opening new opportunities to investigate biological organization across scales.

## Methods

### Ethical statement

All procedures involving animals were reviewed and approved by the responsible governmental authority (Regierung von Unterfranken) and performed in accordance with the German Animal Welfare Act (TierSchG) and relevant institutional guidelines. C57BL/6J mice were housed under a 12 h light/12 h dark cycle at 22–24 °C and 50–60% relative humidity, with ad libitum access to food and water.

### Reagents

Reagents B-27™ Plus Supplement (A3582801), formaldehyde methanol-free (28906), GlutaMAX™ (35050038), 10x HBSS (14060040), Neurobasal™ Plus (A3582901), sodium dodecyl sulfate (SDS, AM9820), Syto16 (S7578), 3-aminopropyl)triethoxysilane (APTES, 440140), Triton™ X-100 (Triton, 28314), trypsin-EDTA (25300054), Tween 20 (28320) were purchased from Thermo Fisher Scientific. Acetic acid (A6283), acrylamide (AA, A4058), ammonium persulfate (APS, A7460), bovine serum albumin (BSA, A7030), B27 Plus supplement (A3582801), N,N′-(1,2-Dihydroxyethylen)bisacrylamid (DHEBA, 294381), N,N-Dimethylacrylamid (DMAA, M1533), dithiothreitol (DTT, 646563), N,N′-methylenebisacrylamide (BA, M1533), ethanol (32205), Gentamycin (G1397), Glutamax (35050061), Neurobasal™ (A3582901), PBS (D8537 and D1408), Penicillin-streptomycin (P4333), poly-D-lysine (PDL, P6407), Sodium Acrylate (408220), Sucrose (S0389) and N,N,N′,N′-tetramethylethylenediamine (TEMED, T7024) were purchased from Merck. For antibodies and NHS-dyes, refer to Supplementary Tables 6-7. GelMap 12mm coverslips with 20 microns patterns of R2-myc-his protein conjugated with ATTO643 were purchased from Visualise.bio.

### Hippocampal neuron preparation

Isolation of primary neurons: E18 C57BL/6J mice were used for primary hippocampal neuron isolation under approval from Bavarian state authorities. Hippocampal tissue underwent digestion in 0.25% trypsin-EDTA at 37°C for 15 min, followed by two washes in a solution of 1x Hanks’ balanced salt solution with Gentamicin. Trituration of hippocampi with varied pipette pore sizes preceded seeding onto poly-D-lysine coated coverslips. Specifically, 40,000 neurons were plated on each 12 mm PDL-coated coverslip (1 mg/mL, 1 h at RT, washed twice with ddH2O). Neurons were cultured in Neurobasal™ medium including 1:50 B27 Plus supplement and 1:100 Glutamax for 21-23 days at 37°C, 5% CO2, with 50% medium replacement weekly.

### Cell lines

COS-7 (monkey kidney) cells were obtained from CLS Cell Line Service GmbH and maintained at 37 °C with 5% CO2 in DMEM supplemented with L-glutamine, 10% FBS, 100 U/mL penicillin, and 0.1 mg/mL streptomycin. For experiments, 75,000 cells per well were seeded onto 12 mm round high-precision coverslips (No. 1.5; Marienfeld, 0117520) placed in 4-well culture plates (Techno Plastic Products, 92012) and cultured for 24 h prior to fixation. HeLa-Nup107-GFP cells were cultured under identical conditions. In this case, 60,000 cells per well were seeded onto the same type of coverslips in 4-well plates and grown for 24 h before fixation. RPE1 (human retina pigment epithelial) cells were seeded and maintained as described previously^69^.

### Murine spermatocyte spread

C57BL/6J mice of 23 and 60 days old were sacrificed using CO_2_, followed by cervical dislocation. Testes were resected and immersed in PBS1X after decapsulation. Next, nuclear spreading’s were carried out as described by^70^, with specific adaptations to improve the results of the expansion microscopy protocols. Briefly, whole decapsulated testes were disrupted with a razor blade, and cells were flushed out by resuspension with a 100 µL pipette. To obtain a cell concentration where it is easier to locate the proper region of the sample, we use 0.5 g of testicular material for 2.5 mL of PBS1X. Afterwards sample was transferred to hypotonic buffer (30 mM Tris-HCl pH 8.2, 17 mM sodium citrate, 5 mM EDTA, 50 mM sucrose, 5 mM DTT) and incubated for 10 min. Then, the hypotonic treatment was stopped by adding an equal volume of sterile 100 mM sucrose. 50 µL of treated cells were dispersed onto a round 12 mm (1.5H) coverslip coated with 1% paraformaldehyde and 0.15% Triton X-100 and allowed to dry overnight in a humid chamber. Coverslips were subsequently stored at -80°C until use.

### Cell culture of Raji B cells

Raji B cells (ATCC, #CCL-86) were cultured in RPMI 1640 (Sigma-Aldrich, #R8758) containing 10% fetal bovine serum, penicillin (100 U/ml), and streptomycin (0.1 mg/ml) at 37 °C and 5% CO_2_. Cells were maintained at a maximum density of ∼2 × 10^6^ cells/ml in standard T25 culture flasks (Sarstedt, #83.3910.502).

### Cell fixation

For neurons, HeLa and COS-7 cells, a fixation solution (FS) composed of 4% formaldehyde + 4% sucrose in 1X PBS was prepared. Fixation was performed at 37°C first for five minutes in a FS solution diluted 1:1 with the media of the cells (to dilute it to 2% FA/Suc), and then full FS was used for 10 minutes, followed by three rinses in PBS1X. RPE1 cells were fixed by cold methanol: the coverslips with cells were plunged into cold methanol (−20 °C) and incubated for 5 min at -20°C as well, followed by three rinses in PBS1X. Raji B cells were incubated for 10 min in PDL-coated coverslips before immersion in the anchoring solution which acted as fixation step as well.

### Iterative expansion

The workflow of Mega-ExM through its three rounds of expansion, as shown in Figure 1a, was performed as follows in this section.

**1. *Anchoring (3 h – overnight)***. Coverslips with sample were incubated in the anchoring solution at 37°C, with different concentrations and time depending on the cell type or main structure of interest. For all experiments in neurons, HeLa, and COS-7 cells (mitochondria and NPC), AS: 2% AA + 1.4% FA in PBS1X for 3 h; spermatocytes (SC) used same AS but overnight. RPE1 cells (centrosomes) were anchored at half this concentration (1% AA; 0.7% FA in PBS1X) but for 5 h.
**2. *1st TREx gel embedding (1.5 h)***. A gelation chamber was built by creating a humid chamber with a petri dish, with a layer Parafilm lying flat on the center. This gelation chamber was put on ice along with the anchored samples. The anchoring solution was discarded and replaced by PBS1X, and the samples and gelation chamber were left for 5min to equilibrate with the ice. Freshly thawed but cold TREx monomer solution (Supplementary Table 1) was supplemented with TEMED/APS to 0.15% final concentration, vortexed, and 30-50uL of gelation solution was placed in the parafilm of the gelation chamber per coverslip. Coverslips were incubated inverted on these drops for 5min on ice and then at 37°C for 1h to allow gelation, along with leftovers of gelation solution as polymerization control.
**3. *Denaturation (1.5 – 2 h)***. After gelation, the coverslip with the gel was carefully removed from the imaging chamber and dipped in 1 mL of denaturation buffer (200 mM Sodium Dodecyl Sulfate (SDS); 200 mM NaCl; 50 mM Tris-BASE) at 37°C for 10min to allow detachment from the coverslip. Then, the gel was transferred to a 1.5 mL centrifuge tube with ∼1.4 mL of fresh denaturation buffer and incubated for 1-1.5h. The pH of the denaturation buffer, the temperature and time of denaturation, also varies along samples. For neurons, HeLa, and COS-7 cells, pH was 6.8, and denaturation was performed for 1.5h at 85°C. For spermatocytes and RPE1 cells, pH was 9.0 and denaturation was done for 1h at 95°C.
**4. *Expansion (2 h)***. After denaturation, gels were left to cool down at RT for ∼10min, before dipping them in petri dishes in PBS1X. Washing of the SDS/denaturation buffer was achieved by exchanging the PBS1X at least 2 times more at intervals of ∼10min. In PBS1X gels expanded ∼2.92x. Then they were cut into smaller pieces and expanded in Type 1 / ultrapure water by 3 washes of 20 min each in the petri dishes.
**5. *Labelling (2 h - 2 days)***. Gels were stained with NHS-dyes first and then immunostained if needed according to the staining section of methods. This order was chosen as first immunostaining and then NHS creates a bias of the NHS staining towards the targeted epitopes, which can be much stronger than the rest of the staining. As we relied on NHS for ultrastructure staining, we performed always NHS first, and we did not find any decrease in the immunostaining quality of all antibodies tested (as there is a possibility of masking epitopes by the NHS staining).
**6. *Neutral gel embedding (1.5 h)*.** Stained expanded gels were checked at the microscope to select regions of interest and good sample density. Then, gels were cut into 10mm x 15mm rectangles, one corner was cut to maintain an asymmetrical shape, and were incubated with neutral gel solution (10% AA + 0.05% DHEBA + 0.1% TEMED + 0.1% APS) under agitation at 4°C for 15 minutes two times in centrifuge tubes or 4-well plates with at least 500 µL of gelation solution per gel. The gelation solution was exchanged for freshly prepared and cold new gelation solution between these incubations. This step allows the gels to fully equilibrate with the neutral gel solution, which shrinks the gel in a controlled manner (less than they shrink in TREx solution if the neutral gel is skipped). After incubation, excess liquid was removed, and the gels were placed inside a humid chamber sandwiched between two round 24 mm (1.5H) coverslips acting as a re-embedding chamber. Polymerization was performed at 37°C for 1 hour. ***Optional***: In the case of antibody labeled samples, after neutral gel embedding, gels were often subjected to a second anchoring step using 1.4% FA and 2% AA solution for at least 3h (up to overnight) at 37°C to secure the antibody-antigen complexes within the expanding matrix, which can slightly improve the signal in the iterative gels but it is not mandatory as the first swellable gel is retained.
**7. *Second TREx embedding (1.5 h).*** Gels were incubated twice for 15 minutes each with TREx monomer solution at 4°C in centrifuge tubes or 4-well plates with at least 500 µL of gelation solution per gel. Following removal of excess liquid, gels were again sandwiched between two round 24 mm (1.5H) within a humid chamber and incubated at 37°C for 1 hour.
**8. *Dissolution of neutral gel (1.5 h).*** After second TREx embedding, gels were incubated in 25mL tubes filled with 200mM NaOH to dissolve the DHEBA-based neutral gel. Then, NaOH was replaced by PBS1X and incubated at least 30 min to neutralize the gel. Gels were later washed two times with Type 1 water, and expansion was achieved by 5h to overnight incubation in at least 1L of water in food-safe 2.5L buckets. After this, gels expanded ∼5x-6x of their original size, i.e., from 10 mm x 15 mm to 50 mm x 75 mm in TNT-ExM stage. Gels were further trimmed xy with rectangular coverslips and razor blades. Gels were usually cut to a height of 2 mm in a petri dish lid by using two pairs of microscope slides spaced ∼ 2 cm as steppers (1 mm height each, stacked in pairs, 2 mm total). The gel was positioned between the two pairs of slides, and a razor blader was slid through the gel using the microscopy glass slides as steppers to fix the 2 mm thickness of the cut slice.
**9. *Mega-*ExM (third round of expansion).** Steps 6, 7, and 8 are a full -NT iteration. For Mega-ExM, as stated in Figure 1, steps 9, 10 and 11 are identical to 6, 7 and 8. We did not found the need of including the optional anchoring step of step 6 in Mega-ExM iteration stages, as the antibodies should have been successfully crosslinked to the second TREx gel, or should survive still attached to the first TREx gel indirectly through the epitopes. For Supplementary Tables 2-3, the rounds of iteration are performed as described as well, no matter the recipes of gels used.

### Post-expansion NHS-dye staining

Expanded gels were equilibrated in NaHCO_3_ 100 mM prepared in PBS1X for 30min. The entire proteome was labeled by incubating the gels with NHS-dyes diluted in the same 100 mM NaHCO_3_ solution. The 10 mm x 15 mm expanded gels shrank to ∼ 3 mm x 5 mm size gels. For neurons, HeLa and COS-7 cells samples, two-three replicate gels of this size were stained together in 125 µL of 80 µg/mL NHS-dye solution in a 0.6 mL centrifuge tube (124 µL of buffer and 1 µL of 10 µg/µL NHS-dye stock dissolved in DMSO). Gels were stained for 1.5h at RT with gentle shaking. For SC (spermatocytes), staining was performed at 80 µg/mL but 1 h RT, and for centrosomes (RPE1 cells), NHS-staining was performed at 40 µg/mL for 1h at RT. Subsequently, gels were transferred to a Petri dish and immersed in ddH2O with 3 exchanges every 30 minutes or until full expansion is achieved. Unless stated otherwise, ATTO643-NHS was used; the rest of the dyes used for the supplementary material, in three or four combinations staining (each individual NHS-dye at 80 µg/mL), is detailed in Supplementary Table 7.

### Post-expansion immunostaining

Expanded NHS-stained gels were equilibrated in 5% BSA solution (prepared in water for SC and PBS1X for the rest) for 30 min. Primary antibodies were applied in 5% BSA solution (same as before) and incubated overnight at 4°C at the dilutions shown in Supplementary Table 6. Gels were washed three times with PBS1X + 0.1% Triton X-100 and then expanded into water once more. Secondary antibodies were applied for 3 hours at 37°C or overnight RT under gentle agitation, according to Supplementary Table 6. Gels were then expanded into water by exchanging the water every 30 minutes until full expansion was achieved (2-3 times).

### Silanization of coverslips

Gels were placed on silanized 1 well chambered cover glass (thickness No. 1.5H, Cellvis C1-1.5H-N) or 24mm cover glass coverslips (thickness No. 1.5H, Marienfeld, 0117640) for imaging. Coverslips or Cellvis were previously cleaned via four sequential sonication in water but for 15 min each first in containers filled with water, then 1M KOH, water again, and finally 99% ethanol in an ultrasound bath for 30 minutes each. After drying, coverslips were incubated with a silane solution consisting of 0.75 % APTES, 90 % ethanol and 5 % acetic acid in ddH2O. Cellvis were incubated with ∼1 mL of silane solution, 24mm round coverslips were submerged in silane solution, both were left under fume hood for 30 minutes. Coverslips and cellvis were washed once with 99 % ethanol for 30 minutes and left to dry. Cellvis and coverslips were stored at -20°C until use.

### Airyscan imaging

Airyscan images were acquired using an LSM 900 microscope equipped with Airyscan 2 (Zeiss) in SR imaging mode. The following lenses from Zeiss were used and are indicated by its corresponding icon in all the images: (i) C-Apochromat 40×/1.2 W Korr; (ii) Plan-Apochromat 20x/0.8 M27, (iii) Plan-Apochromat 10x/0.45 M27, and (iv) EC Plan-Neofluar 2.5x/0.085 M27. Appropriate excitation wavelengths and filter settings for the dyes were chosen through the dye presets in ZEN 2 blue software (Zeiss, version 3.5). All images were processed using the standard strength mode for 3D Airyscan processing.

### Image Analysis Pre/Post-ExM comparison

The expansion factor of the samples was determined by acquiring Airyscan confocal microscopy images of already characterized structures. For spermatocytes SC, we acquired images in TREx, TNT- and Mega-ExM, and we determined the central region’s width from the peak-to-peak distance from the lateral elements, value to which we subtracted (FWHM1 + FWHM2)/2 to have an estimate on the central region, which has been reported to be 117 nm in murine models, which would be the conservative value as others reported it to be 100 nm. Furthermore, we evaluated distortions introduced by the re-embedding process from TREx to TNT with a custom Python script already reported previously by our group^17^. Briefly, corresponding SC in TREx and TNT-ExM regions were acquired with the 40x W lens and 10x Air lens from the Airyscan setup. Then, they were aligned by rigid similarity transformation, and non-rigid affine transformation was then used to calculate deviations from the similarity transformation and generate a distortion vector map. For centrosomes, RPE1 cells were used, and the 200 nm value of the distal was used as ruler for TREx vs TNT-ExM. Similarly, in the multiciliate cells from the hippocampal cultures, 200 nm of distal diameter from the basal body of the cilia was used for TNT-ExM vs Mega-ExM. For mitochondrial cristae from neurons, the average 32 nm was used as explained properly in Figure 5 for TNT-ExM vs Mega-ExM. Furthermore, same features in different stages of the protocol of samples or whole cells’ diameters were compared in Figure 1, Figure 2, and Supplementary Figure 1. For Supplementary Figure 6, cells areas were quantified using the analyzed particles tool of ImageJ in thresholded images.

### Averaging of NPCs in TNT

Center positions of NPC were selected manually in each FOV, and subvolumes were cropped automatically around each particle using a fixed radius and an intensity-dependent height. The extracted subvolumes were centered in the XY plane and then iteratively aligned using log-polar phase correlation of 10-fold upsampled sum projections in all three directions. The rotation angle for the X and Y projections was restricted to +/- 20 degrees to avoid flipping of the images, as the NPC rings were usually parallel to the imaging plane. The normalized sum image of all particles was used as reference for the subsequent iteration. The number of iterations was set to 10, after which the alignment error only improved marginally. For visualization, we additionally applied rotational averaging around the Z axis to each individual aligned particle to impose 8-fold symmetry, if each dye could have bound to the identical site on any of the other 7 subunits of the nuclear pore complex. The alignment and averaging workflow were implemented in python (version 3.12) using scikit-image^71^, scipy^72^, napari^73^ and bioio^74^ packages.

### Determination of NPC size and resolution in TNT

To determine the height and diameter of the expanded NPC, intensity profiles of the sum projections along each axis of the aligned images were computed. Particle boundaries were defined as the position where the intensity falls below 1/3 of the maximum. The NPC ring diameter was taken as the means of the measured values for the XZ and YZ projections. We report both the mean and standard deviation for the individual particles, as well as the size of the symmetrized average particle.

### Resolution estimation using Fourier Shell Correlation (FSC) in TNT

We used “gold standard” FSC to estimate the resolution of the averaged NPC^75–77^. The extracted sub volumes from one FOV were split into two independent half-sets and aligned and reconstructed independently as described above. The two reconstructed average particles were normalized and aligned to each other. A loose soft mask was calculated from the sum of the two NPC and applied to both images. The FSC curve between the soft-masked volumes was determined, and the spatial frequency at which the FSC falls below 0.143 was used as criterion for the resolution estimate.

### Three-dimensional NPC reconstruction from Mega-ExM data

Three-dimensional density maps of nuclear pore complexes (NPCs) were reconstructed from Mega-ExM localization coordinates, as previously described for expansion microscopy-based analysis of individual protein structures^5^. Briefly, Airyscan 3D image stacks from Mega-ExM expanded gels from primary hippocampal neurons were acquired with optimal acquisition for the 40X 1.2 N.A. water lens. Each acquired volume contained up to five individual NPCs, giving a total of 20 NPCs from two different neurons from the same gel. To localize NPCs, air lenses were first used to localize somas from neurons, and gels were trimmed axially to have the base of the nuclei on a reachable distance for the immersion lens. As gels were stained with ATTO643-NHS, individual NPCs were recognized and annotated from (1) their localization in their nuclear membrane, and (2) the presence of three recognizable rings. The resulting 3D image stacks were filtered using a bandpass procedure, followed by an empiric thresholding procedure (eliminating signals under the mean + one standard deviation of the signal in the image), to identify particles (signal spots). To avoid placing duplicate identities on the same particles found in multiple Z sections, the signals were processed using the Crocker and Grier algorithm^76^, and the identities and average coordinates of all particles were determined. Signals clearly originating outside the NPCs were manually removed, and the remaining coordinates were organized into individual NPC datasets for downstream reconstruction. Each input text file represented one NPC and contained x, y and z localization coordinates calibrated to the physical dimensions of the expanded specimen. The first three columns were interpreted as the x, y and z coordinates, respectively. All three spatial dimensions were maintained at the same scale, and no additional axial scaling or correction was applied.

For NPC *i*, we write the coordinate set as

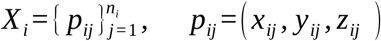

where *p_ij_*denotes localization *j* and *n_i_* is the number of localizations associated with NPC *i*.

Each localization cloud was centred independently by subtracting its mean coordinate:

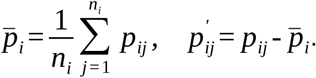

Here, *p_i_* is the mean coordinate vector of NPC *i* and *p^’^* is the centred coordinate of point *j*. This operation removed the absolute position of each NPC within the imaging volume while preserving its internal three-dimensional geometry. Principal component analysis (PCA) was applied to the centred coordinates to align the individual NPCs within a common reference frame. PCA1 was used as the principal reference axis. Following projection into the PCA coordinate system, each localization was represented as

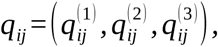

where 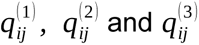 are the components of localization *j* along PCA1, PCA2 and PCA3 respectively.

### Density-map reconstruction and symmetry application

The aligned localization clouds were converted into three-dimensional density volumes by trilinear voxel deposition. The contribution of each localization was distributed fractionally among the neighbouring voxels according to its position within the voxel grid. This cloud-in-cell approach produced a smoother density representation and reduced discretization artefacts compared with nearest-neighbour voxel assignment^77,78^. Two reconstruction conditions were generated from the same localization coordinates. The C1 reconstruction was generated without imposed rotational symmetry. For the C8 reconstruction, eightfold rotational symmetry was imposed around the PCA-defined reference axis. Full-data reconstructions were used for visualization. For Fourier shell correlation (FSC) analysis, the 20 original NPCs were divided into two non-overlapping groups before symmetry application. Each group was reconstructed independently to generate two half-maps. For the C8 condition, eightfold symmetry was applied independently to each half-map, thereby maintaining the independence of the particle subsets used for resolution estimation. The independent sample size remained defined by the 20 original NPC localization files and was not increased by symmetry application. An isotropic voxel size of 750 nm in expanded-sample coordinates was used for density-map reconstruction. This value was chosen to balance coordinate sampling density and preservation of local structural detail, given that the reconstruction was based on 20 separate NPC localization datasets. After correction using the measured expansion factor of 215, this corresponds to approximately 3.49 nm in the native biological scale.

### Expansion factor estimation

The expansion factor was determined from the 20 individual NPC localization files. A native NPC reference diameter of 100 nm was used for scale calibration. As PCA1 defined the reference axis, the perpendicular distance of each localization from this axis was calculated from its PCA2 and PCA3 components:

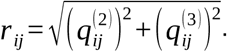

The root mean square (RMS) radius of NPC *i* was calculated as

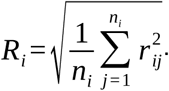

The expanded NPC diameter *D_i_* and the corresponding expansion factor *E F_i_* were then calculated as

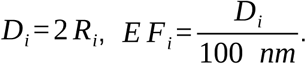

Across the 20 coordinate files, the mean expanded RMS diameter is 21.528 ±0.374 µm, reported as mean ± SEM. This corresponds to a mean expansion factor of 215 ± 3.74 (mean ± SEM). For FSC reporting, we rounded this value to EF = 215.

### Fourier shell correlation analysis for NPCs in Mega-ExM

The resolution of the reconstructed density maps was estimated from independently generated half-maps using version 3.0 of the 3DFSC program suite^79^. FSC analysis was restricted to reconstructions generated using a voxel size of 750 nm in expanded-sample coordinates, corresponding to approximately 3.49 nm in the native biological scale after expansion correction with EF = 215. The spatial-frequency axis was converted to the native biological scale using the experimentally determined expansion factor of 215. Resolution was reported using the FSC 0.143 criterion^80,81^. The C1 and C8 reconstructions were evaluated independently. After expansion correction using EF = 215, the FSC 0.143 resolutions were 3.46 nm (34.62 Å) for the C1 reconstruction and 3.31 nm (33.05 Å) for the C8 reconstruction. The higher consistency of the C8 reconstruction reflects the imposed rotational symmetry and does not represent an increase in the number of independent NPC particles.

### Data presentation and statistical analysis

For all data that was not from NPC, plots were made in OriginPro 2023b software (OriginLab Corporation, Northampton, MA, USA.) When distributions of measurements passed a Shapiro-Wilk normality test, the mean ± standard error of the mean (SEM) was used to describe the population. If data was not normal e.g. skewed, the median ± median absolute deviation (MAD) was used additionally to the mean ± SEM. For NPC data, refer to the respective section in methods.

## Acknowledgements

The authors thank E. Maier, I. Simeonov, and L. Behringer-Pließ for cell culture and technical support; Manfred Alsheimer (Cell and Developmental Biology -Zoology I-Department, JMU Wuerzburg) for supplying mouse testis; Nadine Kraft, Claudia Groh, and Wolfgang Rössler (Behavioral Physiology and Sociology -Zoology II-Department, JMU Wuerzburg) for supplying bee brain tissue; as well as Bettina Warscheid and Lakshita Sharma (Biochemistry II Department, JMU Wuerzburg) for supplying the OXPHOS antibody cocktail. I.V.V, C.W., A. H.S., S.O.R., and M.S. acknowledge funding from the European Research Council (ERC) under the European Union’s Horizon 2020 research and innovation program (grant agreement No 835102). I.V.V and C.G-N. were funded by scholarships from the German Academic Exchange Service (DAAD), “Doctoral Programmes in Germany, 2023/24 (57645448) and 2025/26 (57730835), respectively. O.I.G.M acknowledges funding from Agencia Nacional de investigación e Innovación (ANII, Uruguay), CSIC grupos I+D 2022 grant (to A. Geisinger and R.B.) and Programa de Desarrollo de las Ciencias Básicas (PEDECIBA). The authors gratefully acknowledge the computing time granted by the Resource Allocation Board and provided on the supercomputer Emmy/Grete at NHR-Nord@Göttingen as part of the NHR infrastructure. The calculations for this research were conducted with computing resources under the project nib00040. This work was also supported by DFG project SFB1286/A03 to S.O.R.

## Author contributions

I.V-V and O.I.G.M developed the concept and designed experiments. I.V-V, performed and analyzed ExM of hippocampal neurons cultures, HeLa, COS-7 cells and Raji cells. O.I.G.M performed and analyzed ExM experiments from spermatocytes. C.G-N. performed experiments from RPE1 cells. G.W. performed ExM experiments for expansion chemistry. C.W. prepared hippocampal mouse neurons. P.E. prepared Raji cells cultures. J.T.H. prepared the supplementary movies. For the Mega-ExM NPC data analysis and reconstruction: S.O.R. carried out the preprocessing step and generated the x, y and z coordinates required for the 3D reconstruction; A.H.S. devised the 3D reconstruction strategy; A.H.S. and S.O.R supervised the development of the 3D reconstruction pipeline; S.C. developed the 3D reconstruction pipeline and generated the 3D structure; C.G. performed the 3DFSC analysis; and S.C. supervised the 3DFSC calculation. G.P. supplied the RPE1 cell cultures and centrosome-related antibodies. S.O.R. performed the analysis for Mega NPCs. P.K. performed the analysis on TNT NPCs. R.B supervised the study. M. S. conceived the concept, designed experiments and supervised the study. I.V-V and M.S. wrote the manuscript. All authors reviewed and approved the manuscript.

## Data availability

Image data are available from the corresponding authors on reasonable request.

## Supplementary information

Supplementary material is available.

## References

1. Balzarotti F, Eilers Y, Gwosch KC. Nanometer resolution imaging and tracking of fluorescent molecules with minimal photon fluxes. Science. 2016;355:606–612.

2. Reinhardt SCM, Masullo LA, Baudrexel I. Ångström-resolution fluorescence microscopy. Nature. 2023;617:711–716.

3. Sahl SJ, Matthias J, Inamdar K. Direct optical measurement of intramolecular distances with angstrom precision. Science. 2024;386:180–187.

4. Chen F, Tillberg PW, S BE. Expansion Microscopy. Science. 2015;347:543–548.

5. Shaib AH, Chouaib AA, Chowdhury R. One-step nanoscale expansion microscopy reveals individual protein shapes. Nat Biotechnol. 2025;43:1539–1547.

6. Taban D, Jungblut M, Budiarta M. Single-step expansion SMLM enables molecular-resolution imaging in cells and isolated proteins. Chem Biomed Imaging. 2026;4:1530–1537.

7. Baumeister W. Cryo-electron tomography: A long journey to the inner space of cells. Cell. 2022;185:2649–2652.

8. Young LN, Villa E. Bringing structure to cell biology with cryo-electron tomography. Annu Rev Biophys. 2023;52:573–595.

9. Helmerich DA, Beliu G, Taban D. Photoswitching fingerprint analysis bypasses the 10-nm resolution barrier. Nat Methods. 2022;19:986–994.

10. Helmerich DA, Sauer M. Challenges and Limitations of Molecular Resolution Fluorescence Imaging, Methods Appl. Fluoresc. 2025;13:043101.

11. Helmerich DA, Budiarta M, Eiring P. Impact of Docking Strand Design on Spatial Resolution in DNA Points Accumulation for Imaging in Nanoscale Topography. ChemPhysChem. 2026;27:202500803.

12. Tillberg P, Chen F, Piatkevich K. Protein-retention expansion microscopy of cells and tissues labeled using standard fluorescent proteins and antibodies. Nat Biotechnol. 2016;34:987–992.

13. Ku T, Swaney J, Park JY. Multiplexed and scalable super-resolution imaging of three-dimensional protein localization in size-adjustable tissues. Nat Biotechnol. 2016;34:973–981.

14. Eilts J, Jungblut M, A. HD. Resolving protein organization in cells with nanometer resolution. Nat Commun. 2026;17:7583.

15. Valdes PA, Yu CC (Jay), Aronson J, et al. Improved immunostaining of nanostructures and cells in human brain specimens through expansion-mediated protein decrowding. Sci Transl Med. 2024;16(732):eabo0049. doi:10.1126/scitranslmed.abo0049

16. M’Saad O, Bewersdorf J. Light microscopy of proteins in their ultrastructural context. Nat Commun. 2020;11(1):3850. doi:10.1038/s41467-020-17523-8

17. M’Saad O, Cairns A, Gulcicek J. All-optical visualization of specific molecules in the ultrastructural context of brain tissue. Nat Biotechnol. Published online 2025. doi:10.1038/s41587-025-02905-4.

18. Chang JB, Chen F, Yoon YG. Iterative expansion microscopy. Nat Methods. 2017;14:593–599.

19. Truckenbrodt S, Sommer C, Rizzoli SO, Danzl JG. A practical guide to optimization in X10 expansion microscopy. Nat Protoc. 2019;14:832–863.

20. Damstra HGJ, Mohar B, Eddison M. Visualizing cellular and tissue ultrastructure using Ten-fold Robust Expansion Microscopy (TREx. eLife. 2022;11:73775.

21. Louvel V, Haase R, Mercey O. iU-ExM: nanoscopy of organelles and tissues with iterative ultrastructure expansion microscopy. Nat Commun. 2023;14:7893.

22. Tavakoli MR, Lyudchik J, Januszewski M, et al. Light-microscopy-based connectomic reconstruction of mammalian brain tissue. Nature. 2025;642(8067):398–410. doi:10.1038/s41586-025-08985-1

23. Passmore JB, Serweta AK, Kapitein LC. Beyond 4×: pathways to higher expansion factors in expansion microscopy. Methods Microsc. Published online 2026. doi:10.1515/mim-2025-0040

24. Zwettler FU, Reinhard S, Gambarotto D. Molecular resolution imaging by post-labeling expansion single-molecule localization microscopy (Ex-SMLM. Nat Commun. 2020;11:3388.

25. Shi X, Li Q, Dai Z. Label-retention expansion microscopy. J Cell Biol. 2021;220:202105067.

26. Sarkar D, Kang J, Wassie AT. Revealing nanostructures in brain tissue via protein decrowding by iterative expansion microscopy. Nat Biomed Eng. 2022;6:1057–1073.

27. Hu H, Krah D, Ntolkeras A, et al. *Thousandfold Expansion Microscopy*. the Preprint Server for Biology; 2026.

28. Gambarotto D, Zwettler FU, Le Guennec M. Imaging cellular ultrastructures using expansion microscopy (U-ExM. Nat Methods. 2019;16:71–74.

29. Klimas A, Gallagher BR, Wijesekara P. Magnify is a universal molecular anchoring strategy for expansion microscopy. Nat Biotechnol. 2023;41:858–869.

30. Damstra HG, Passmore JB, Serweta AK, et al. GelMap: intrinsic calibration and deformation mapping for expansion microscopy. Nat Methods. 2023;20:1573–1580.

31. Page SL, Hawley RS. The genetics and molecular biology of the synaptonemal complex. Annu Rev Cell Dev Biol. 2004;20:525–558.

32. Fraune J, Schramm S, Alsheimer M, Benavente R. The mammalian synaptonemal complex: protein components, assembly and role in meiotic recombination. Exp Cell Res. 2012;318:1340–1346.

33. Schücker K, Holm T, Franke C, Sauer M, Benavente R. Elucidation of synaptonemal complex organization by super-resolution imaging with isotropic resolution. Proc Natl Acad Sci USA. 2015;112:2029–2033.

34. Wang Y. Combined expansion microscopy with structured illumination microscopy for analyzing protein complexes. Nat Protoc. 2018;13:1869–1895.

35. Zwettler FU, Spindler MC, Reinhard S. Tracking down the molecular architecture of the synaptonemal complex by expansion microscopy. Nat Commun. 2020;11:3222.

36. Hamel V. Identification of chlamydomonas central core centriolar proteins reveals a role for human WDR90 in ciliogenesis. Curr Biol. 2017;27:2486–2498.

37. Guichard P, Laporte MH, Hamel V. The centriolar tubulin code. Semin. Cell Dev Biol. 2023;137:16–25.

38. C. GS, V AK. The primary cilium: a signalling centre during vertebrate development. Nat Rev Genet. 2010;11:331–344.

39. O’Callaghan C, Sikand K, Rutman A. Respiratory and Brain Ependymal Ciliary Function. Pediatr Res. 1999;46:704.

40. Gray MW. Mitochondrial Evolution. Cold Spring Harb Perspect Biol. 2012;4:011403.

41. McArthur K, Ryan MT. Resolving mitochondrial cristae: introducing a new model into the fold. EMBO J. 2020;39:105714.

42. Davies KM, Strauss M, Daum B. Macromolecular organization of ATP synthase and complex I in whole mitochondria. Proc Natl Acad Sci USA. 2011;108:14121–14126,.

43. Frey TG, Perkins GA, Ellisman MH. Electron tomography of membrane-bound cellular organelles. Annu Rev Biophys Biomol Struct. 2006;35:199–224.

44. Wang C, Østergaard L, Hasselholt S, Sporring J. A semi-automatic method for extracting mitochondrial cristae characteristics from 3D focused ion beam scanning electron microscopy data. Commun Biol. 2024;7:377.

45. Eilts J, Reinhard S, Michetschläger N, Werner C, Sauer M. Enhanced synaptic protein visualization by multicolor super-resolution expansion microscopy. Neurophotonics. 2023;10(4):044412. doi:10.1117/1.NPh.10.4.044412

46. S. C, A. EJ, L S. Mitochondrial Cristae: Where Beauty Meets Functionality. Trends Biochem Sci. 2016;41:261–273.

47. Huang C, Deng K, Wu M. Mitochondrial cristae in health and disease. Int J Biol Macromol. 2023;235:123755.

48. Gu J, Wu M, Guo R. The architecture of the mammalian respirasome. Nature. 2016;537:639–643.

49. Mühleip A, Flygaard RK, Baradaran R. Structural basis of mitochondrial membrane bending by the I–II–III2–IV2 supercomplex. Nature. 2023;615:934–938.

50. Zheng W, Chai P, Zhu J. High-resolution in situ structures of mammalian respiratory supercomplexes. Nature. 2024;631:232–239.

51. Sazanov L. A giant molecular proton pump: structure and mechanism of respiratory complex I. Nat Rev Mol Cell Biol. 2015;16:375–388.

52. Letts J, Fiedorczuk K, Sazanov L. The architecture of respiratory supercomplexes. Nature. 2016;537:644–648.

53. Waltz F, Righetto RD, Lamm L. In-cell architecture of the mitochondrial respiratory chain. Science. 2025;387,1296–1301.

54. Beck M, Hurt E. The nuclear pore complex: understanding its function through structural insight. Nat Rev Mol Cell Biol. 2017;18:73–89.

55. Allegretti M, Zimmerli CE, Rantos V. In-cell architecture of the nuclear pore and snapshots of its turnover. Nature. 2020;586:796–800.

56. Hinshaw JE, Carragher BO, Milligan RA. Architecture and design of the nuclear pore complex. Cell. 1992;69:1133–1141.

57. Appen A. In situ structural analysis of the human nuclear pore complex. Nature. 2015;526:140–143.

58. Stoffler D, Feja B, Fahrenkrog B, Walz J, Typke D, Aebi U. Cryo-electron tomography provides novel insights into nuclear pore architecture: implications for nucleocytoplasmic transport. J Mol Biol. 2003;328(1):119–130.

59. Maimon T, Elad N, Dahan I, Medalia O. The Human Nuclear Pore Complex as Revealed by Cryo-Electron Tomography. Structure. 2012;20:998–1006.

60. Mahamid J, Pfeffer S, Schaffer M. Visualizing the molecular sociology at the HeLa cell nuclear periphery. Science. 2016;351:969–972.

61. Morgan KJ, Carley E, Coyne AN. Visualizing nuclear pore complex plasticity with pan-expansion microscopy. J Cell Biol. 2025;224:202409120.

62. Tan Y, Baldwin P, Davis J. Addressing preferred specimen orientation in single-particle cryo-EM through tilting. Nat Methods. 2017;14:793–796.

63. Verbeke EJ, Gilles MA, Bendory T. Self-Fourier shell correlation: properties and application to cryo-ET. Commun Biol. 2024;7:101.

64. Beck M, Baumeister W. Cryo-electron tomography: Can it reveal the molecular sociology of cells in atomic detail? Trends Cell Biol. 2016;26:825–837.

65. Ledbetter M, Porter KR. A “microtubule” in plant cell fine structure. J Cell Biol. 1963;19:239–250.

66. Wen G, Lycas MD, Yuqing J. Trifunctional Linkers Enable Improved Visualization of Actin by Expansion Microscopy. ACS Nano. 2023;17:20589–20600.

67. Johnson LV, Walsh ML, Bockus BJ, Chen LB. Monitoring of relative mitochondrial membrane potential in living cells by fluorescence microscopy. J Cell Biol. 1981;88:526–535.

68. Sheard TMD, Shakespeare TB, Seehra RS. Differential labelling of human sub-cellular compartments with fluorescent dye esters and expansion microscopy. Nanoscale. 2023;15:18489–18499.

69. Streubel JMS, Karasu OR. Nek1 defines a branch of centriolar microtubule length control parallel to CP110-Cep97. Nat Commun. 2026;17:7330.

70. Peters AHFM, Plug AW, Vugt MJ, Boer P. A drying-down technique for the spreading of mammalian meiocytes from the male and female germline. Chromosome Res. 1997;5(1):66–68.

71. Walt S, Schönberger JL, Nunez-Iglesias J. scikit-image: Image processing in Python. PeerJ. 2014;2:453.

72. Virtanen P, Gommers R, Oliphant TE. SciPy 1.0: fundamental algorithms for scientific computing in Python. Nat Methods. 2020;17:261–272.

73. Sofroniew N, Lambert T, Bokota G. napari: a multi-dimensional image viewer for Python. *Zenodo*. Published online 2022.

74. Brown EM, Toloudis D, Sherman J, et al. BioIO: Image Reading, Metadata Conversion, and Image Writing for Microscopy Images in Pure Python. Computer software]. GitHub; 2023. https://github.com/bioio-devs/bioio

75. Scheres SH, Chen S. Prevention of overfitting in cryo-EM structure determination. Nat Methods. 2012;9:853–854.

76. Crocker JC, Grier DG. Methods of Digital Video Microscopy for Colloidal Studies. J Colloid Interface Sci. 1996;179:298–310.

77. Hockney RWE, E JW. Computer Simulation Using Particles. CRC Press; 1988. doi:10.1201/9780367806934.

78. Canelhas DR, Stoyanov T, Lilienthal AJ. A Survey of Voxel Interpolation Methods and an Evaluation of Their Impact on Volumetric Map-Based Visual Odometry. In: 2018 IEEE International Conference on Robotics and Automation (ICRA. 2018:3637–3643. doi:10.1109/ICRA.2018.8461227.

79. Tan YZ, Baldwin PR, Davis JH, et al. Addressing preferred specimen orientation in single-particle cryo-EM through tilting. Nat Methods. 2017;14:793–796.

80. Bracewell RN. Fourier Analysis and Imaging. 2003. doi:10.1007/978-1-4419-8963-5.

81. Heel M, Schatz M. Fourier shell correlation threshold criteria. J Struct Biol. 2005;151:250–262.

