## Supplementary Information for "Structural cell biology by mega-expansion microscopy"

### I. Supplementary Tables

**Supplementary Table 1. Composition of ExM recipes tested.** We hypothesized that an iterative ExM workflow where only the neutral gel is cleaved and the first swellable gel is retained should be possible. This would maximize the expansion as the neutral gel would not restrict the second swellable gel, while the retention of the first gel would improve overall signal retention. For this purpose, we explored TREx[1], ProExM[2], U-ExM[3], Magnify[4], Cloro(B)[5], and MS1[6] gel recipes and tested their ability to be iterated (continues in Supplementary Table 2).

| Recipe | AA | DMAA | BA | DHEBA | SA | NaCl | PBS | APS | TEMED | Total Monomer | Reference |
|---|---|---|---|---|---|---|---|---|---|---|---|
| TREx | 14.25% | - | 0.009% | - | 10.34% | - | 1X | 0.15% | 0.15% | 24.60% | Damstra, H. G., et al., 2022 |
| ProExM | 2.50% | - | 0.15% | - | 8.60% | 11.70% | 1X | 0.20% | 0.20% | 11.25% | Tillberg, P. W., et al., 2016 |
| U-ExM | 10.00% | - | 0.10% | - | 19.00% | - | 1X | 0.50% | 0.50% | 29.10% | Gambarotto, D., et al., 2018 |
| Magnify | 10.00% | 4.00% | 0.01% | - | 34.00% | 1.00% | 1X | 0.25% | 0.04% | 48.01% | Klimas, A., et al., 2023 |
| Chloro(B) | 10.80% | - | 0.04% | - | 30.10% | - | 1.1X | 0.10% | 0.10% | 40.94% | Bos, P. R., et al., 2024 |
| MS1 | 10.00% | - | - | 0.10% | 19.00% | - | - | 0.25% | 0.25% | 29.10% | Louvel, V., et al., 2023 |

AA=Acrylamide, DMAA=N,N-Dimethylacrylamide, BA=N,N'-methylenebisacrylamide (Bis-Acrylamide), DHEBA=N,N′-(1,2-Dihydroxy-1,2-ethanediyl)bis(acrylamide), SA=Sodium Acrylate. All percentages are expressed as the final concentration of each component in the gelation solution.

**Supplementary Table 2. Iterative ExM combinations tested.** We tested all ExM protocols listed in Supplementary Table 1, both, iterated with itself and with TREx. We chose a TREx-based approach to maximize the expansion factor, while keeping the gelation and re-embedding as robust as possible, as other protocols were highly prone to premature gelation, which renders the experiment unreliable. Even when reducing the APS/TEMED to 0.1% (as done for example in iU-ExM[6] which would be MS1 as 1st gel and U-ExM[3] as 2nd gel), ProExM[2], U-ExM[3] and Magnify[4] gelated prematurely while being re-embedded when used as 2nd gel. Interestingly, when PNP, UNU, and CNC are successfully created and the neutral gel cleaved, the second swellable gel fails to expand the first gel. UNT and CNT also fail to expand beyond the first gel.

| Workflow | 1st Gel Recipe | 2nd Gel Recipe | ExF 1st Gel (PBS1X) | ExF 1st Gel (Water) | Shrinkage NeutralGel | Re-embedding 2nd Gel*** | ExF (final) |
|---|---|---|---|---|---|---|---|
| TNT* | TREx | TREx | 2.83x | 8.15x | 70-80% | Successful | 43.20x |
| PNP | ProExM | ProExM | 2.06x | 4.30x | 75-85% | Premature gelation | - |
| UNU | U-ExM | U-ExM | 2.30x | 4.50x | 70-80% | Premature gelation | - |
| MNM** | Magnify | Magnify | 3.00x | 6.20x | 70-80% | Premature gelation | - |
| CNC | Chloro(B) | Chloro(B) | 2.60x | 5.05x | 70-80% | Premature gelation | - |
| iTREx | MS1 | TREx | 2.58x | 4.40x | 85-90% | Successful | 24.75x |
| PNT | ProExM | TREx | 2.06x | 4.30x | 75-85% | Successful | 18.50x |
| UNT | U-ExM | TREx | 2.30x | 4.50x | 70-80% | Successful | Does not expand |
| MNT** | Magnify | TREx | 3.00x | 6.20x | 70-80% | Premature gelation | - |
| CNT | Chloro(B) | TREx | 2.60x | 5.05x | 70-80% | Successful | Does not expand |

ExF = Expansion Factor. * = if neutral gel is not cleavable (10% AA + 0.025% BA + 0.05% TEMED + 0.05% APS, from LICONN[7]), TNT reaches ~20x expansion. ** = When the first gel is not empty, and methacrolein is used, success rate increases and they reach ~40X (as TNT). *** Except for TREx, all gels used 0.1% APS/TEMED. All ExF are an average of two independent gels iterated simultaneously for each condition, and for each gel the expansion of the width and length of 10 mm x 15 mm starting gels was averaged.

**Supplementary Table 3. Multiple rounds of TREx re-embedding.** We then tested how many rounds of re-embedding (NT), neutral gel dissolution and expansion were possible with this combination in empty gels for TREx, ProExM and MS1, and found that three additional rounds of TREx embedding and expansion were possible, which reached a cumulative expansion factor (ExF) of >2,000x when starting with TREx (TNTNTNT, where Mega is TNTNT, and Giga TNTNTNT). We validated the biological internal expansion factor of three rounds (TREx, TNT, Mega) by acquiring the same HeLa cell in the unexpanded state and through all these rounds, which yielded ~7x, ~40X, ~230X and is shown in Figure 1b-e (for the cell itself, not the gel). A fourth round was possible, which exhibited 5.6x additional expansion when measuring the gel itself, which would imply a maximum expansion factor of ~1,300x for the HeLa cell, but we were unable to confirm this by imaging as the signal was too diluted to be detected. Similarly, the Mega gel of Figure 3g was further iterated, which already demonstrated ~42x for TNT and ~273x for Mega according to the measurement of the basal body diameter (Figure 4f). This gel did expand ~6.0x more, which would imply a potential ~1600x expansion, but we were again unable to acquire any image due to signal intensity dilution. For this reason, Raji B cells gels were stained sequentially two times one hour RT with 80µg/mL of ATTO643-NHS, which was barely enough to detect cells at this stage, but sufficient to estimate the expansion factor by the area of the cell (see Supplementary Figure 6).

| Starting Gel | 1st round ExF | 2nd round ExF | 3rd round ExF | 4th round ExF |
|---|---|---|---|---|
| TREx | 8.15x | 5.3x (43.2x) | 7.5x (320x) | 7.25x (2320x) |
| ProExM | 4.30x | 4.3x (18.5x) | 5.0x (92.25x) | 6.5x (600x) |
| MS1 | 4.40x | 5.5x (24.75x) | 4.5x (111.38x) | 7x (777x) |
| TREx (Fig1b-e, HeLa cell) | ~7.0x | ~5.8x (~40x) | ~5.7x (~228x) | *5.6x (1276x) |
| TREx (Fig3g, Hipp. culture) | ~7.0x | ~6.0x (~42x) | ~6.5x (~273x) | *6.0x (1638x) |
| TREx (SuppFig. 6, Raji cells) | ~5.84x | ~5.00x (~29.0x) | ~8.60x ~(249x) | 6.17x (1532x) |

For the first three rows, all ExFs are an average of two independent gels iterated simultaneously for each condition. For each gel the expansion of the width and length of 10 mm x 15 mm starting gels was averaged. From the second round on, the cumulative expansion factor is shown in parentheses.

**Supplementary Table 4. Statistical values of NPC particles from different datasets.** Average diameter of individual particles and from the reconstruction in the rings and the distances between rings, of TNT expanded NHS-stained samples, from different types of cells. Individual datasets (rows) are from a single field of view (ROI) of a single nucleus (Cell) of the specified cell type (Type). Hippocampal neurons and HeLa cells were analyzed from two independent experiments (Rep), and only one from COS-7, but at least two nuclei per cell type. All values are presented in microns, except for the adjusted FSC values which are in nanometers.

| Type-Rep-Cell-ROI | ROIs | avg height | avg diameter | height of avg | diameter of avg | FSC_no-sym | FSC_sym | ExF | FSC_no-sym_adj | FSC_sym_adj |
|---|---|---|---|---|---|---|---|---|---|---|
| Neurons-1-1-1 | 13 | 7.4 | 4.35 | 7.28 | 4.88 | 1395 | 870 | 39.5 | 35.3 | 22.0 |
| Neurons-1-1-2 | 11 | 7.5 | 3.9 | 7.87 | 4.03 | 1638 | 989 | 35.5 | 46.2 | 27.9 |
| Neurons-1-2-1 | 12 | 7.06 | 4.73 | 7.21 | 5.2 | 1315 | 697 | 43.0 | 30.6 | 16.2 |
| Neurons-1-2-2 | 11 | 6.63 | 3.75 | 6.83 | 3.97 | 1654 | 868 | 34.1 | 48.5 | 25.5 |
| Neurons-2-1-1 | 18 | 7.03 | 4.05 | 7.02 | 4.36 | 1127 | 689 | 36.8 | 30.6 | 18.7 |
| Neurons-2-1-1 | 19 | 7.19 | 4.2 | 7.02 | 4.75 | 1134 | 865 | 38.2 | 29.7 | 22.7 |
| HeLa-1-1 | 35 | 4.08 | 2.95 | 4.09 | 3.12 | 785 | 628 | 26.8 | 29.3 | 23.4 |
| HeLa-1-2 | 55 | 4.74 | 2.31 | 4.88 | 2.41 | 752 | 513 | 21.0 | 35.8 | 24.4 |
| HeLa-2-1 | 40 | 4.67 | 2.51 | 4.75 | 2.79 | 454 | 337 | 22.8 | 19.9 | 14.8 |
| HeLa-2-2 | 32 | 4.01 | 2.18 | 4.09 | 2.27 | 651 | 557 | 19.8 | 32.8 | 28.1 |
| COS7-1-1 | 26 | 4.66 | 2.81 | 4.81 | 3.06 | 1091 | 716 | 25.5 | 42.7 | 28.0 |
| COS7-1-2 | 27 | 5.13 | 2.54 | 5.33 | 2.73 | 861 | 646 | 23.1 | 37.3 | 28.0 |
| **Average Neurons** | **84** | **7.14** | **4.16** | **7.21** | **4.53** | **1377** | **830** | **37.8** | **36.4** | **21.9** |
| **Average HeLa** | **162** | **4.38** | **2.49** | **4.45** | **2.65** | **661** | **509** | **22.6** | **29.2** | **22.5** |
| **Average COS-7** | **53** | **4.90** | **2.68** | **5.07** | **2.90** | **976** | **681** | **24.3** | **40.1** | **28.0** |

**Supplementary Table 5. List of NHS-dyes esters used.**

| Dye | Catalog | Vendor |
|---|---|---|
| Atto488 | AD 488-301 | Leica Microsystems |
| Atto550 | AD550-31 | Leica Microsystems |
| Atto633 | AD633-31 | Leica Microsystems |
| Atto643 | AD643-31 | Leica Microsystems |
| Atto647N | AD647N-35 | Leica Microsystems |
| Atto665 | AD655-31 | Leica Microsystems |
| CF555 | 92130 | Biotium |
| CF405M | BOT-92111 | BIOZOL |
| Cy3 | 21020 | Lumiprobe |
| CF568 | SCJ4600027 | Sigma-Aldrich |
| Alexa Fluor 488 | A20000 | ThermoFisher |
| Fluorescein | 46410 | ThermoFisher |
| Pacific Blue | P10163 | ThermoFisher |

**Supplementary Table 6. Summary on 13 different NHS-dyes staining in TREx stage of MegaExM protocol.** We chose to characterize the NHS-dyes according to how enriched is the signal in the six different compartments/organelles that can be easily recognized by morphology in 8-fold expanded TREx when HeLa cells were crosslinked with 2% AA + 1.4% FA 3h 37°C, and gels were denatured at 85°C pH=6.8 (MegaExM protocol; examples shown through Supplementary Figures 6,7,8,9). "/ " = not evaluated, as CF405 was quite dim when stain in competition with other NHS-dyes, and no mitotic HeLa with enough signal was found. "-" = negative staining, as it is not stained or cannot be recognized by NHS alone. "+", "++", "+++" means positive staining of the compartment, where +++ is the strongest signal in the field of view, and fainter signals are ++ and +.

| NHS-dye | Nucleolus | Nucleoplasm | NPC | Mitochondria | Centrosome | Chromosomes |
|---|---|---|---|---|---|---|
| CF405 | +++ | + | + | - | / | / |
| Pacific Blue | +++ | ++ | ++ | ++ | ++ | + |
| Fluorescein | +++ | + | ++ | - | +++ | + |
| Atto488 | +++ | ++ | + | + | +++ | ++ |
| Alexa488 | +++ | ++ | + | ++ | +++ | ++ |
| Atto550 | +++ | + | +++ | +++ | +++ | + |
| CF555 | +++ | ++ | ++ | ++ | +++ | ++ |
| CF568 | +++ | ++ | + | - | ++ | ++ |
| Cy3 | +++ | ++ | ++ | +++ | +++ | - |
| Atto633 | +++ | + | +++ | ++ | +++ | + |
| Atto643 | ++ | ++ | ++ | +++ | +++ | - |
| Atto647N | ++ | ++ | ++ | +++ | ++ | ++ |
| Ato665 | +++ | ++ | ++ | +++ | +++ | - |

**Supplementary Table 7. List of antibodies used in the present work.**

| Type | Host | Target(Conjugation) | Dilution | Catalogue | Vendor/Reference |
|---|---|---|---|---|---|
| Primary | Mouse | Acetylated-tubulin | 1:100 | Homemade | Streubel, J. M. S., et al., 2026 |
| Primary | Mouse | Acetylated-tubulin | 1:250 | 66200-1-Ig | Proteintech |
| Primary | Mouse | OXPHOS | 1:250 | 45-8199 | ThermoFisher |
| Primary | Rabbit | Alfa-tubulin | 1:500 | 11224-1-AP | Proteintech |
| Primary | Rabbit | CEP83 | 1:500 | HPA038161 | Sigma-Aldrich |
| Primary | Rabbit | CEP164 | 1:500 | Homemade | Schmidt, K. N., et al., 2012 |
| Primary | Rabbit | Homer1 | 1:100 | 160003 | Synaptic Systems |
| Primary | Rabbit | ODF2 | 1:100 | Homemade | Kuhns, S., et al., 2013. |
| Primary | Rabbit | SYCP3 | 1:500 | Homemade | Baier, A., et al., 2007 |
| Primary | Rabbit | vGlut1 | 1:100 | 135 303 | Synaptic Systems |
| Primary | Guinea Pig | vGlut1 | 1:100 | 135 318 | Synaptic Systems |
| Secondary, F(ab')2 | Goat | Mouse IgG (Alexa Fluor™488) | 1:250 | A11017 | ThermoFisher |
| Secondary, F(ab')2 | Goat | Rabbit IgG (Alexa Fluor™555+) | 1:250 | A48283 | ThermoFisher |
| Secondary, F(ab')2 | Goat | Rabbit IgG (CF™568) | 1:250 | SAB4600310 | Sigma-Aldrich |
| Secondary, F(ab')2 | Goat | Guinea Pig IgG (Alexa Fluor™488) | 1:250 | JIM-106-546-003 | Jackson ImmunoResearch |

### II. Supplementary Figures

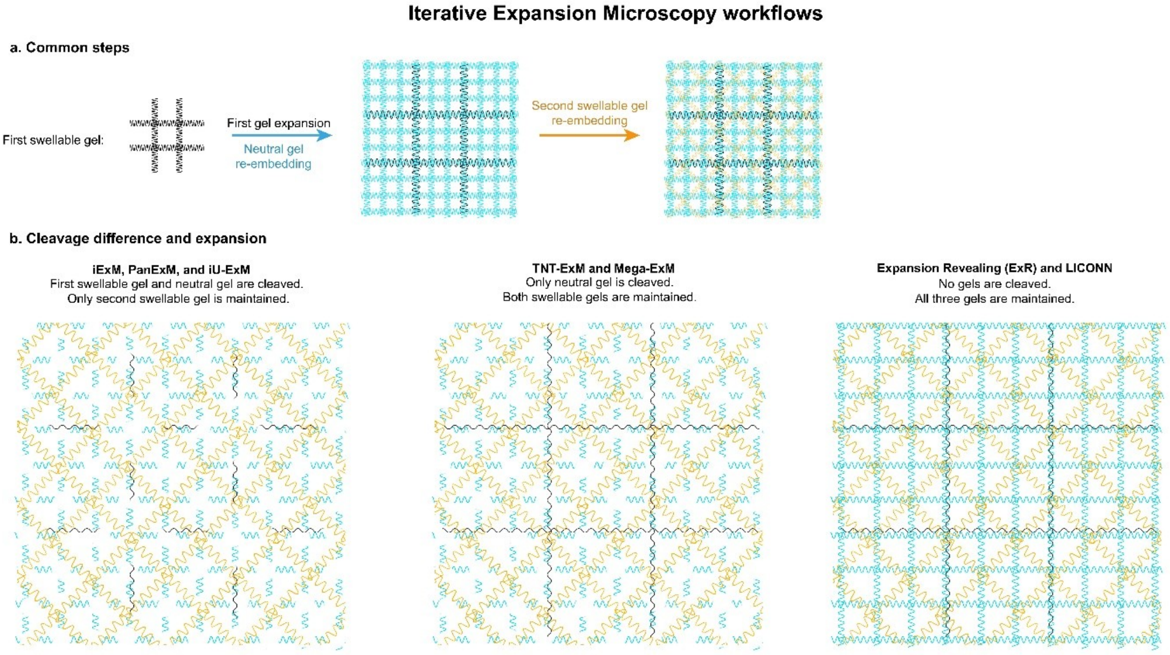


**Supplementary Fig. 1. Overview on iterative expansion microscopy methods. a**, Published iterative ExM methods are based on the original iExM workflow, where a swellable gel is stabilized by a neutral non-swellable gel before embedding in another swellable gel[8]. Distinct protocols diverge on which gel is cleaved to ensure expansion, if a cleavage step is done at all. **b**, Scheme showing the first classical approaches using DHEBA as the crosslinker of the first swellable gel and the neutral gel, resulting in a cleavage of all but the last swellable gel (left side). A more recent approach shown first in expansion revealing (ExR)[9] and later by LICONN[7], avoids any cleavage at all as it uses bis-acrylamide as the crosslinker of all the gels (b, right). We developed TNT-ExM and Mega-ExM as an in-between approach, cleaving only the neutral gel to maximize the expansion factor (without cleaving it, TNT reaches ~20X instead of ~40-50X), but retaining all the swellable gel to maximize the signal (b, middle).

a

| Gel # | SA (M) | AA (M) |
|---|---|---|
| 1 | 0.275 | 2.825 |
| 2 | 0.55 | 2.55 |
| 3 (TREx) | 1.1 | 2.1 |
| 4 | 1.6 | 1.5 |
| 5 | 2.2 | 0.9 |
| 6 | 3.0 | 0.1 |
| 7 | 3.1 | 0 |

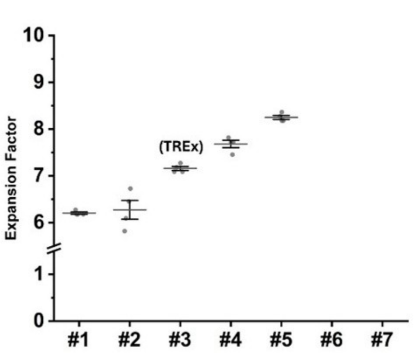


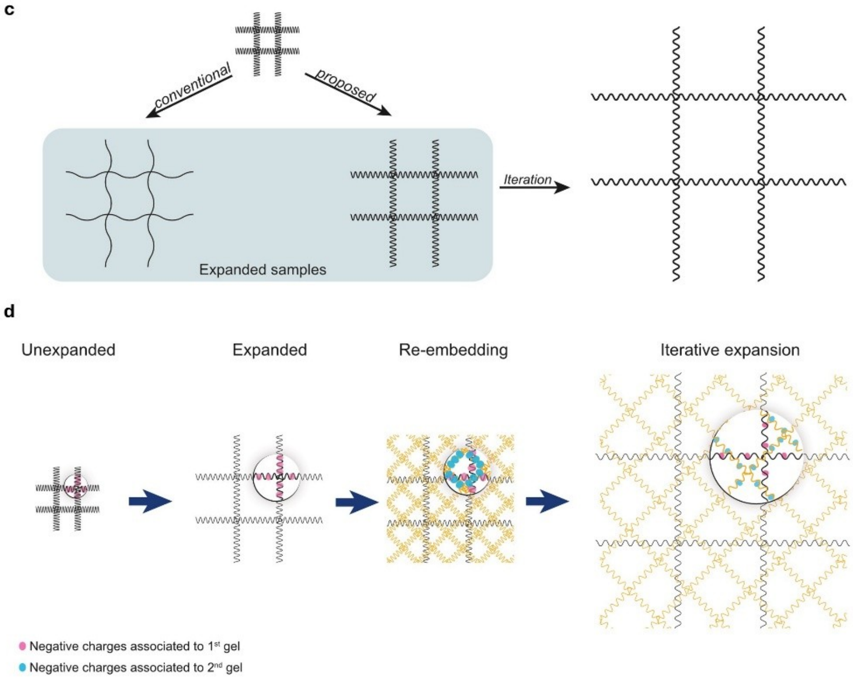


**Supplementary Fig. 2. Expansion of the gel is limited by the amount of acrylate that it contains. a**, Previous attempts on achieving higher expansion factors have been mainly focused on decreasing the amount of the bis-acrylamide crosslinker, which comes with a compromise in gel structural integrity. Here, we explored the effect of the acrylate itself on the TREx recipe, while keeping the bis-acrylamide unchanged. We explored seven combinations of sodium acrylate (SA) and acrylamide (AA), while keeping its combined amount unchanged to keep the overall monomer concentration. The specific concentrations are shown in table (a). **b**, Expansion factors achieved by different combinations of sodium acrylate (SA) and acrylamide (AA), with recipes #6 and #7 missing as the gelation of those recipes were not successful. Recipe #3 is TREx as published. **c**, We hypothesize that gels achieve their expansion plateau because the negative charges of acrylate are separated enough to reach an equilibrium in which their repulsion forces are no longer sufficient to further expand the gel (c, proposed), not because the mesh of the gel has reached a physical limit (c, conventional) as it is usually portrayed. The proposed model explains why gels can further expand in the second and third swellable gel in MegaExM where only neutral gel is cleaved, or other protocols where no gel is cleaved[7]. **d**, Scheme showing the distribution of negative acrylate charges of the first and second gel, and how they allow for further expansion. In Mega, three TREx gels are present at the same time in the final gel.

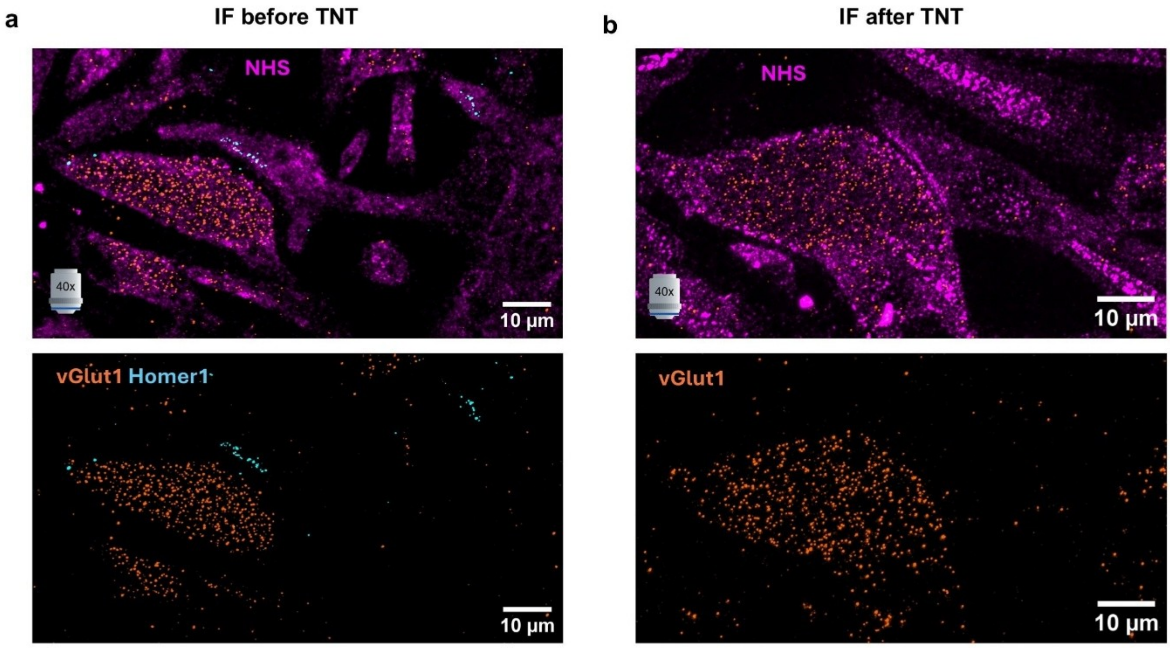


**Supplementary Fig. 3. Immunolabeling after one and two expansion rounds**. **a,b**, Airyscan images of dye-NHS and immunolabeled vGlut1 (pre-synaptic) and Homer1 (post-synaptic) in primary hippocampal neurons. Labeling was performed after the first (TREx) (a) and second (TNT) (b) expansion round. After TNT expansion only vGlut1 was imunostained. As we did not observe any differences in image quality, we proceeded with immunostaining at the TREx stage in all other experiments.

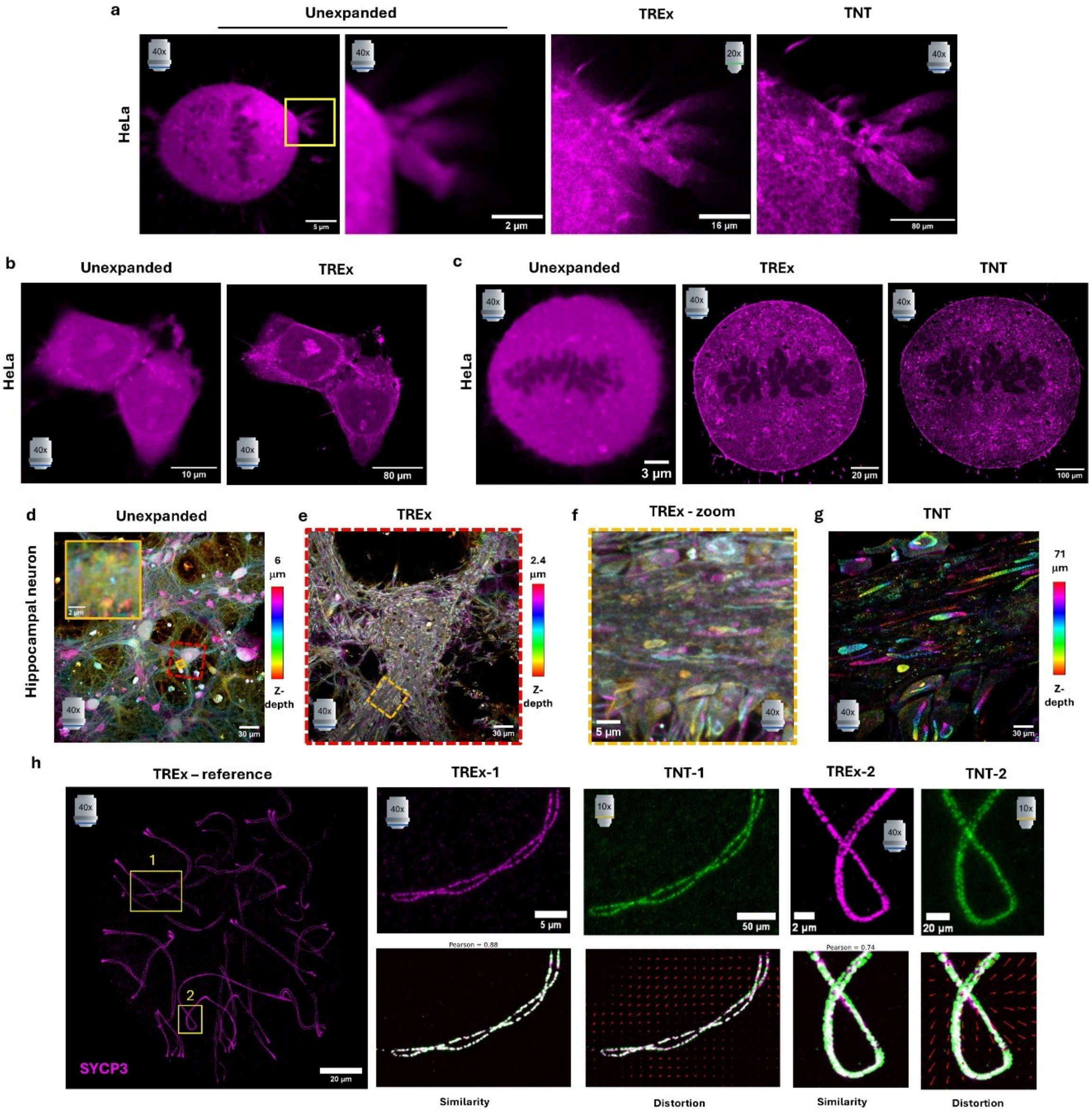


**Supplementary Fig. 4. Pre/post expansion comparison across unexpanded, TREx and TNT stages. a**, Unexpanded mitotic HeLa cell and zoomed-in in one appendix, which is further depicted in TREx and TNT for size comparison. Similarly, (b) shows two HeLa cells between unexpanded and TREx, and a whole mitotic HeLa cell (c) throughout TREx and TNT. Furthermore, (d) shows a hippocampal culture which is further expanded across TREx and TNT, with focus on the pyramidal neuron enclosed in the red rectangle, which is then showed in (e) in TREx. Furthermore, the continuous yellow rectangle from (d) and the discontinuous yellow rectangle from (e) highlight the same primary dendrite of this neuron, which is showed in (f) as a zoomed-in region of (e) and its correlative acquisition in TNT in (g). d-g are shown as color-coded z-projections. Images of the same synaptonemal complexes from mouse spermatocytes are shown in (h) to compare TREx with TNT. First with the whole nuclei projection as reference in the TREx stage, and then two synaptonemal complexes indicated by the yellow rectangles are compared between their TREx and TNT expanded states. The synaptonemal complexes were registered by similarity transformation, yielding a Pearson correlation coefficient between the two images and a distortion vector map showing the differences between the similarity and affine transformation. a-g correspond to ATTO643-NHS stained samples, and (h) shows the post-expansion immunostaining against SYCP3.

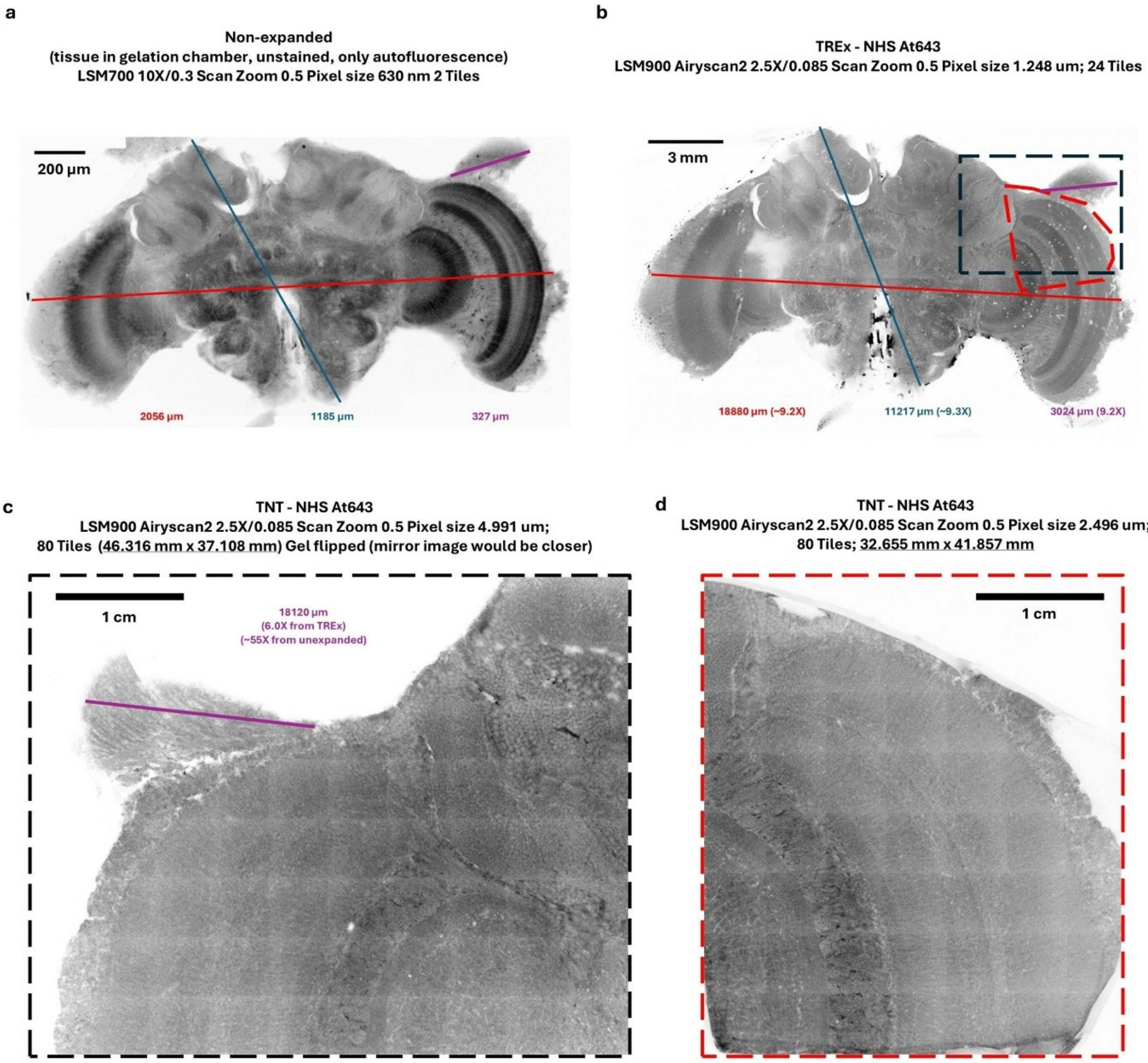


**Supplementary Fig. 5. TNT-ExM on brain tissue.** To evaluate if tissues can be further expanded/iterated by our protocol, we performed TNT in honeybee (*Apis mellifera*) workers. Slices were prepared as described in Kraft, N., et al., 2023 and then proceed to MegaExM workflow. After gelation, still inside the gelation chamber, the tissue was scanned in a LSM700 confocal setup to have a reference on the starting size **(a).** Then, after the first TREx, which was denaturated at 95°C for 100min in the buffer pH=9.0, the tissue was stained with ATTO643-NHS and expanded, achieving approximately 9-fold expansion when measuring the dimensions of the tissue compared to PreExM **(b vs a)**. One hemisphere of the brain was cut and further iterated, starting as a square gel of ~1.4 cm width, which in TNT reached over the 7 cm width. As this did not fit the imaging chamber the gel was blindly cut and scanned, with the result being the image shown in **(c)**. With a recognizable feature of the tissue, we estimated the internal expansion factor (purple line in **b** and **c**) as at least 50X fold (55X from the measurement), which agrees to the ~5x expansion of the gel (1.4 cm to 7 cm). The gel was further trimmed cleanly and result image is shown in **d**.

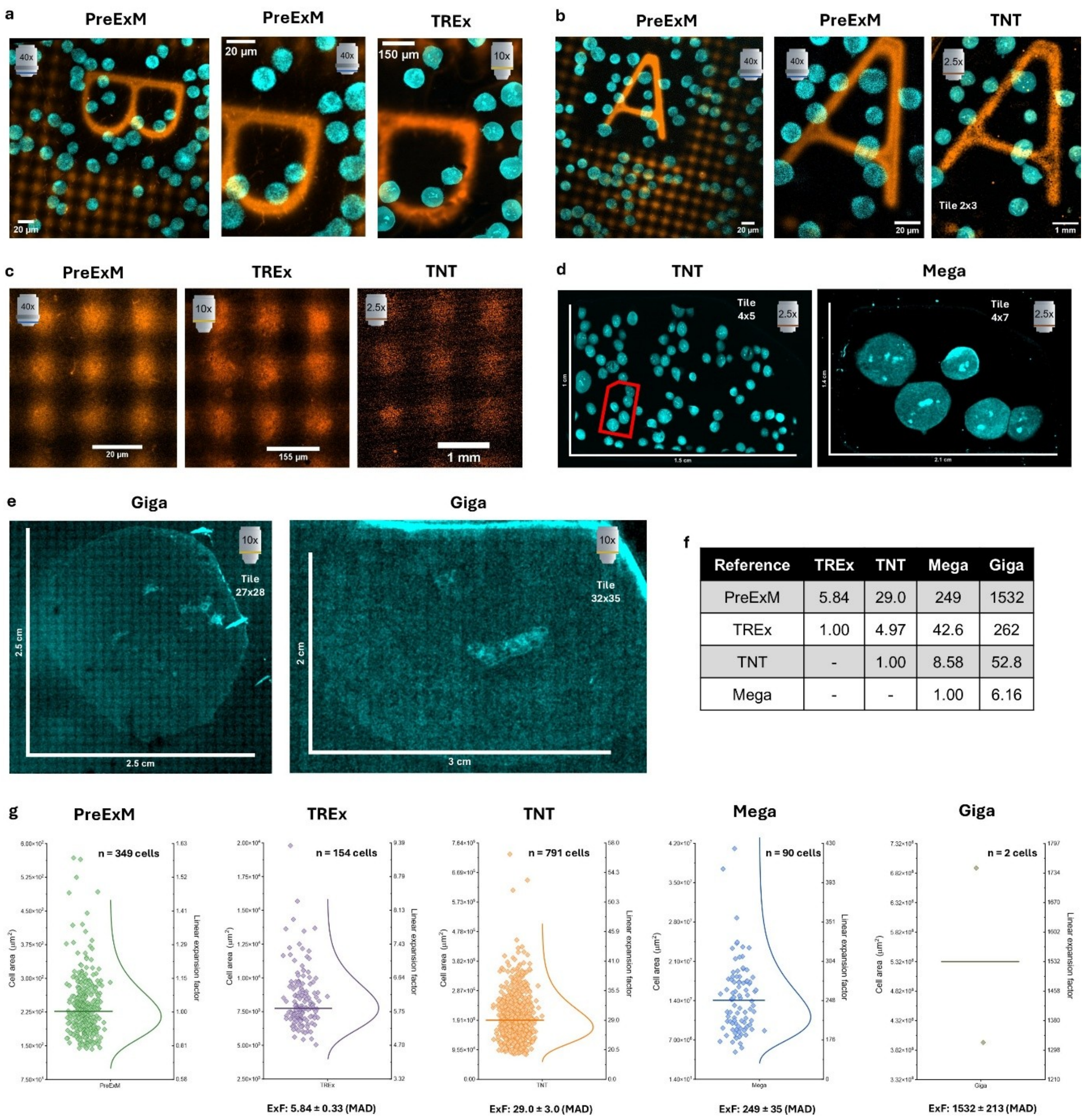


| Reference | TREx | TNT | Mega | Giga |
|---|---|---|---|---|
| PreExM | 5.84 | 29.0 | 249 | 1532 |
| TREx | 1.00 | 4.97 | 42.6 | 262 |
| TNT | - | 1.00 | 8.58 | 52.8 |
| Mega | - | - | 1.00 | 6.16 |

**Supplementary Fig. 6. Validation of four rounds of expansion with B cells plated in GelMap grid coverslips.** Raji B cells were seeded on GelMap patterned coverslips and subjected to Mega-ExM. **a**,**b**, Comparison of TREx and PreExM cells/grid (with unstained cells, only autofluorescence) (a) while (b) compares TNT with PreExM. **c,** Expansion of the 20 microns pattern of GelMap through TREx and TNT. **d,** As GelMap did not give sufficient signal after a third round of expansion, a TNT gel of 1 cm x 1.5 cm with Raji B cells is shown (left), which was subsequently expanded to Mega and further trimmed into a 1.4 cm x 2.1 cm gel size (right). The corresponding area of the Mega gel in the TNT gel is highlighted in red, which had an original size of ~2 mm x 3.3 mm. **e**, Giga gels were produced by subjecting Mega gels to a fourth round of expansion, which successfully expand ~6x from Mega surpassing ~1500x linear expansion (≥ 2 cm diameter). **f,** Comparison of the linear expansion factor of all four rounds compared to PreExM by the measured cell area (f). **g,** Distribution of all cells sizes measured in the five stages of the protocol. Data is always shown as Median ± Median Absolute Deviation (MAD).

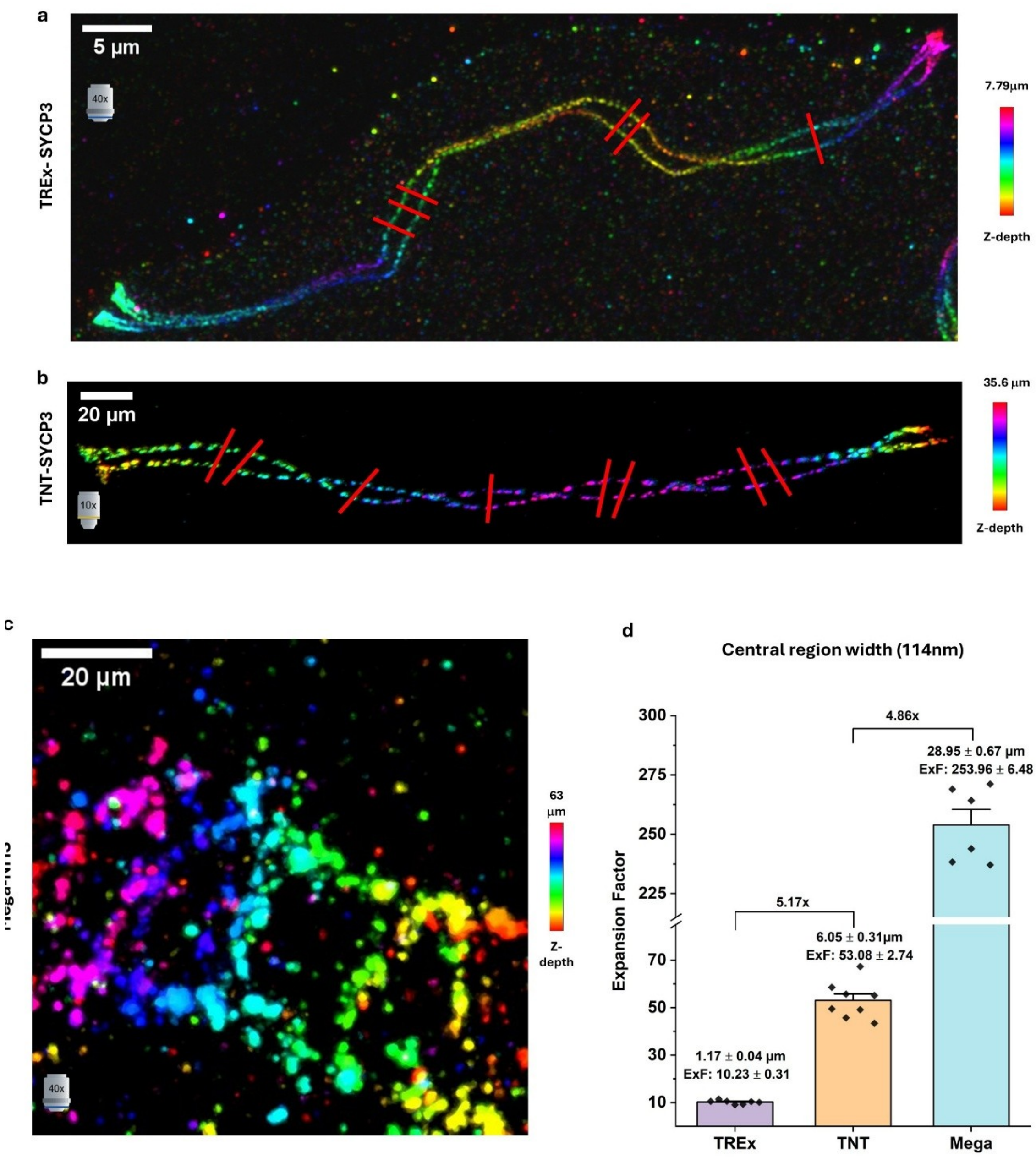


**Supplementary Fig. 7. Synaptonemal complexes (SCs) as nanoruler**. **a,b**, Airyscan images of TREx- and TNT-expanded SCs immunolabeled for lateral elements SYCP3 and with ATTO643-NHS, respectively. The expansion factor was determined by measuring the distance between the lateral filaments at the widest points using perpendicular line profiles. **c**, Airyscan image of a ATTO643-NHS labelled SC. **d**, Average expansion factors achieved. The central region width was estimated by subtracting the average FWHM of both peaks to the peak-to-peak distance of the lateral elements (d), which yielded ~10X, ~50X, and ~250X when using the reported 117 nm width for murine synaptonemal complex[8,9]. Mean values measured for the central region were 1.17 ± 0.04 μm for TREx, 6.05 ± 0.31 μm for TNT, and 28.95 ± 0.67 μm for Mega (Mean ± S.E.), for one SC per stage of the protocol (7 line-profiles for TREx, 8 for TNT, 6 for Mega).

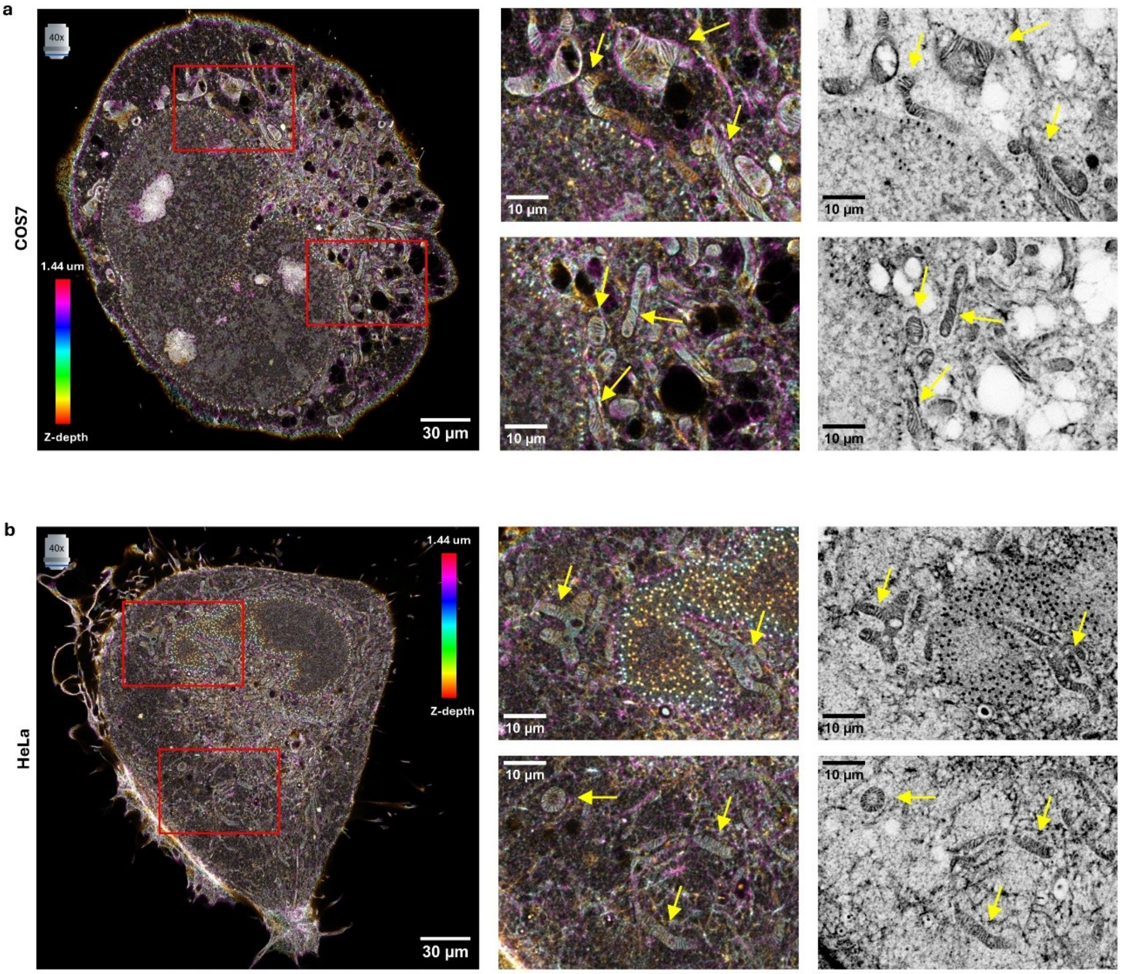


**Supplementary Fig. 8. Mitochondrial cristae of TREx-expanded COS-7 and HeLa cells**. As mitochondrial inter-cristae distance from cell lines like COS-7 (a) and HeLa (b) are nearly thrice as big than hippocampal neurons (~80-100 nm compared to ~30 nm), TREx is enough to resolve this mitochondrial feature. **a,b**, Airyscan images of ATTO643-NHS-stained gels are shown with a focus on mitochondria, first as z-color coded projections and then inverted grayscale on the right. Red rectangles show the areas amplified, and yellow arrows highlight individual mitochondria with cristae resolved.

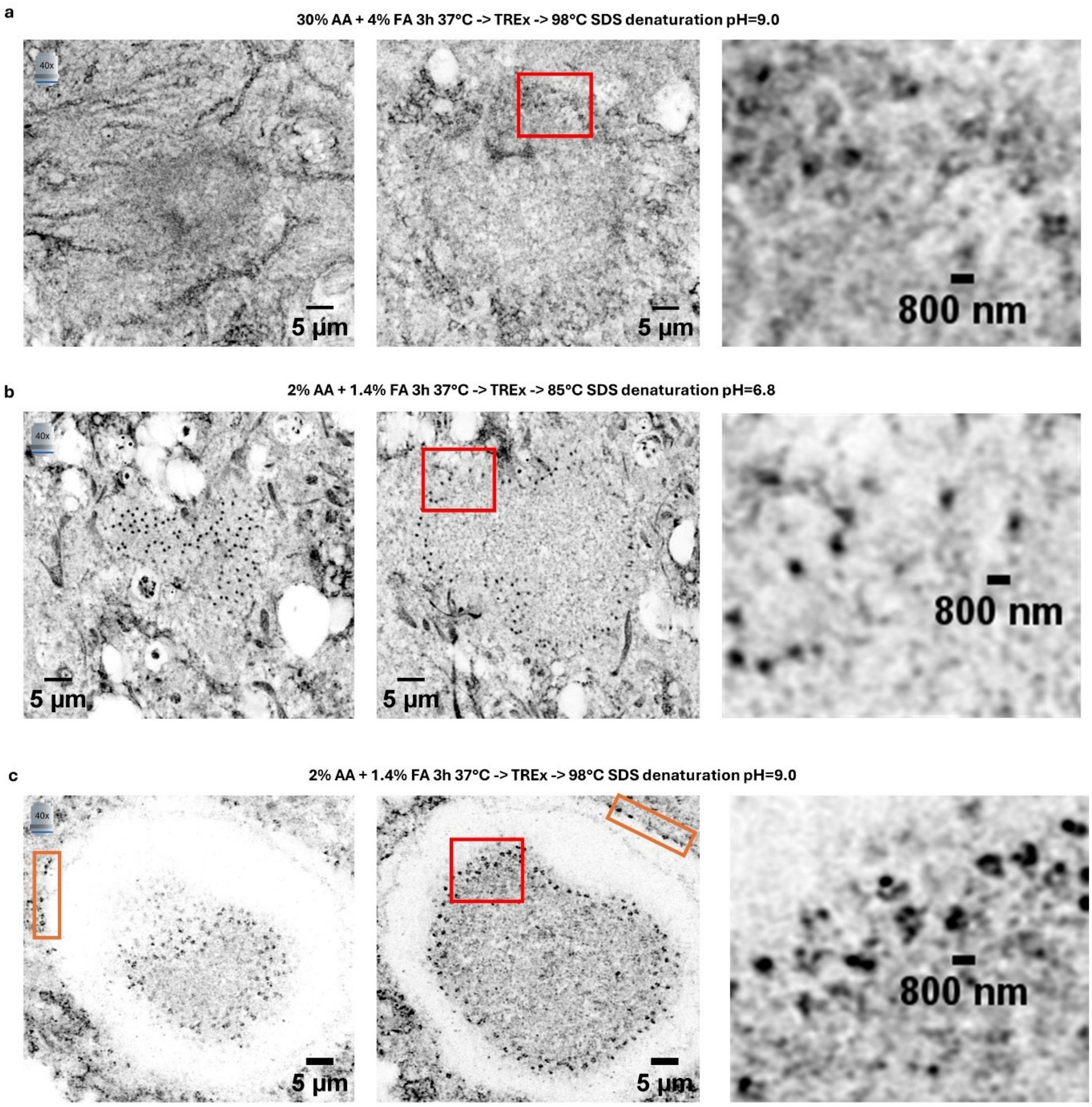


**Supplementary Fig. 9**. **Optimizing anchoring/crosslinking and denaturation Mega-ExM of NPCs.** We studied different combinations of Acrylamide (AA) + formaldehyde (FA) crosslinking/anchoring with common denaturation conditions, and its effect on the NHS staining pattern. **a**, First, we combined the MAP protocol[10] (30% AA/4% FA for anchoring and 98°C, pH=9.0 for denaturation) with TREx gelation of primary hippocampal cultures, which resulted in NHS patterns that were not highly enriched in any compartment, but was sufficient to recognize NPCs in the nuclei base as ~600-800 nm diameter rings or dots. **b**, Next, we tried the anchoring and denaturation from iU-ExM[6], which based on U-ExM[3] and Pan-ExM[11] using lower amounts of AA/FA but an in-between denaturation temperature and lower pH to avoid the disintegration of the first cleavable gel while maintaining a good labeling and expansion of centrioles (85°C pH=6.8 instead of 73° or pH=9.0). This protocol provided the best results and was used for the Mega-ExM workflow. **c**, In addition, the protocol (b) avoids the decoupling of the nuclei from the cytoplasm during expansion, which happens in almost all the cells when using lower anchoring (2% AA + 1.4% FA) but stronger denaturation (98°C and pH=9.0). Decoupling of the nuclei tends to happen along the NPCs, as usually the cytoplasmatic and nuclear rings are separated, and the first one is retained in the cytoplasm (orange rectangles) while the nuclear ones are kept in the nuclei (red rectangle).

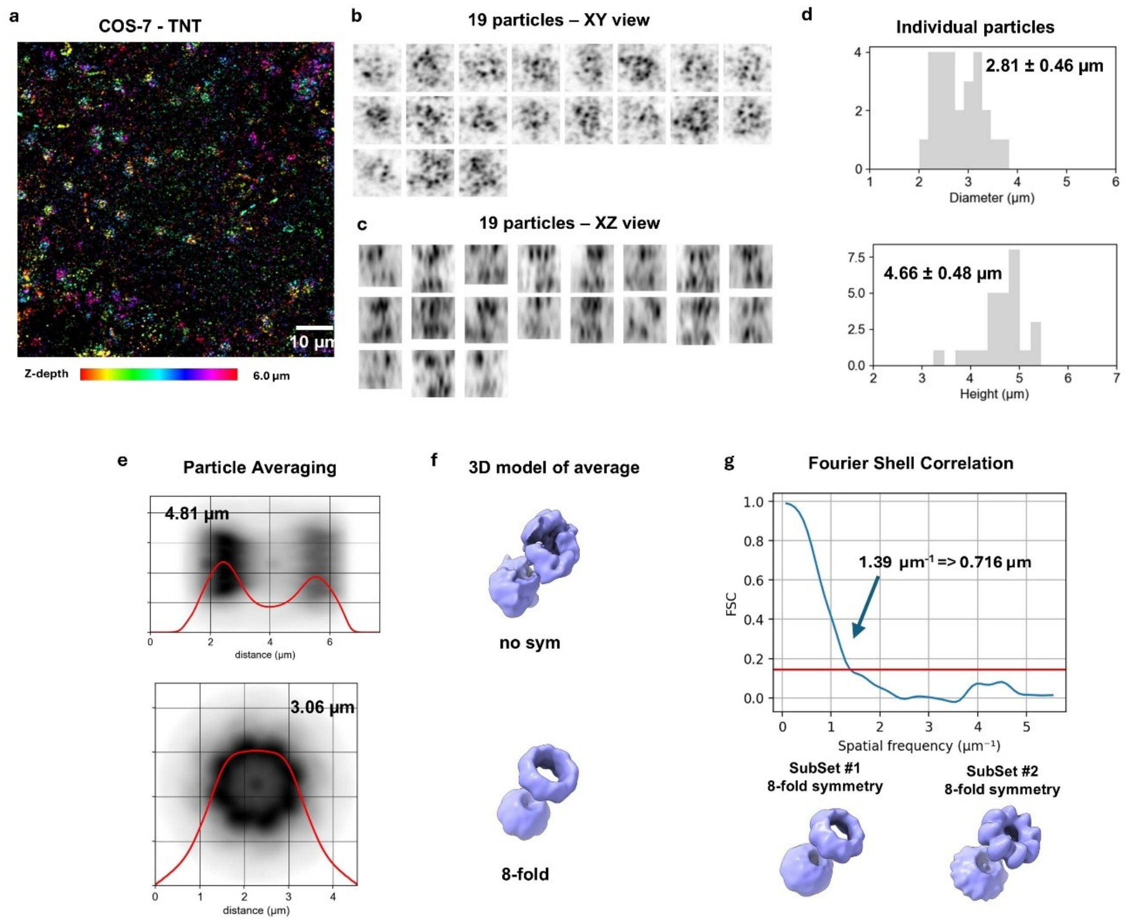


**Supplementary Fig. 10. NPC reconstruction by particle averaging of ATTO643-NHS labeled TNT-expanded COS-7 cells**. **a,** 3D-Airyscan image of a TNT-expanded COS-7 cell, focusing on the bottom of the nuclei, shown as z-color-coded projection, where NPC can be visualized as rings. **b,c,** Example isolated particles (individual NPCs) shown in their frontal *xy* view (b) and side *yz* view in (c), where at least two rings (cytoplasmic and nuclear) can be recognized. **d**, Histograms of ring diameters and distances between rings (height) detected for individual NPCs. **e**, Reconstruction of an average TNT-expanded NPC using Fourier correlation alignment of N=19 particles for a single field of view of one nucleus of COS-7 cell, with 8-fold symmetry imposed, with intensity profiles (red) superimposed. **f**, 3D reconstruction models of averaged particles with and without 8-fold symmetry imposed. **g**, Estimation of practical resolution in the reconstruction of the shown data set by Fourier shell correlation[64,65] of two datasets of NPCs acquired from the same gel and same COS-7. All values included are from the dataset identified as COS7-1-1 in the first column of Supplementary Table 4. Values shown as Mean +/- SD.

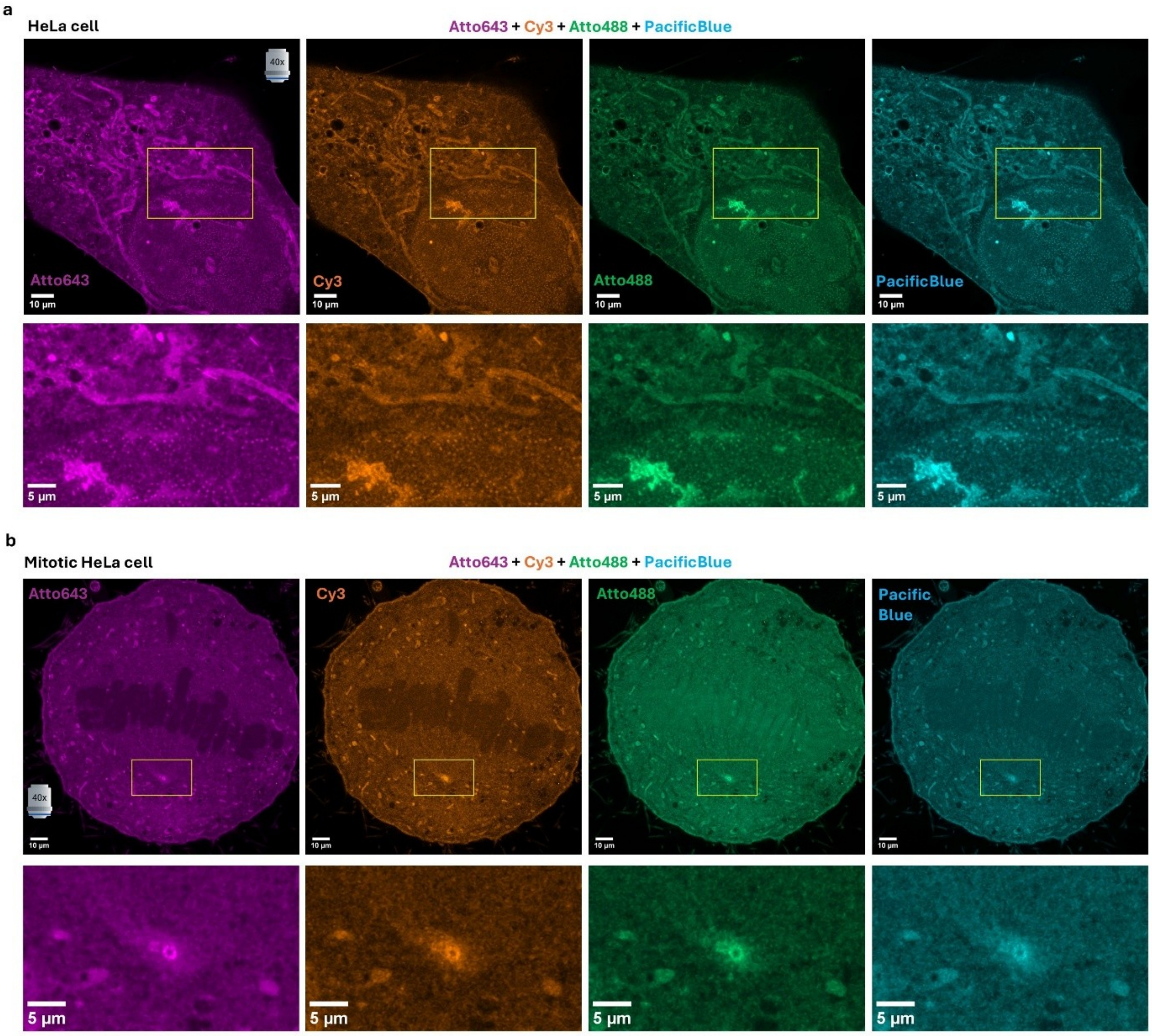


**Supplementary Fig. 11**. **TREx-expanded cells stained with different NHS-functionalized dyes (part 1 of 4)**. To evaluate if different NHS-dyes have any bias on their staining towards certain compartments of the cells, as they should have different affinities due to the variable partitioning coefficient of the dye structure (hydrophobic vs hydrophilic), we stained gels of HeLa cells with the combination of ATTO643-NHS + Cy3-NHS + ATTO488-NHS + PacificBlue-NHS. **a**, HeLa cell in G0, with focus on the nuclear pores and mitochondria (yellow rectangle, zoomed below). **b**, Mitotic HeLa cell, with focus on the centrosome (yellow rectangle, zoomed below) and the chromosomes.

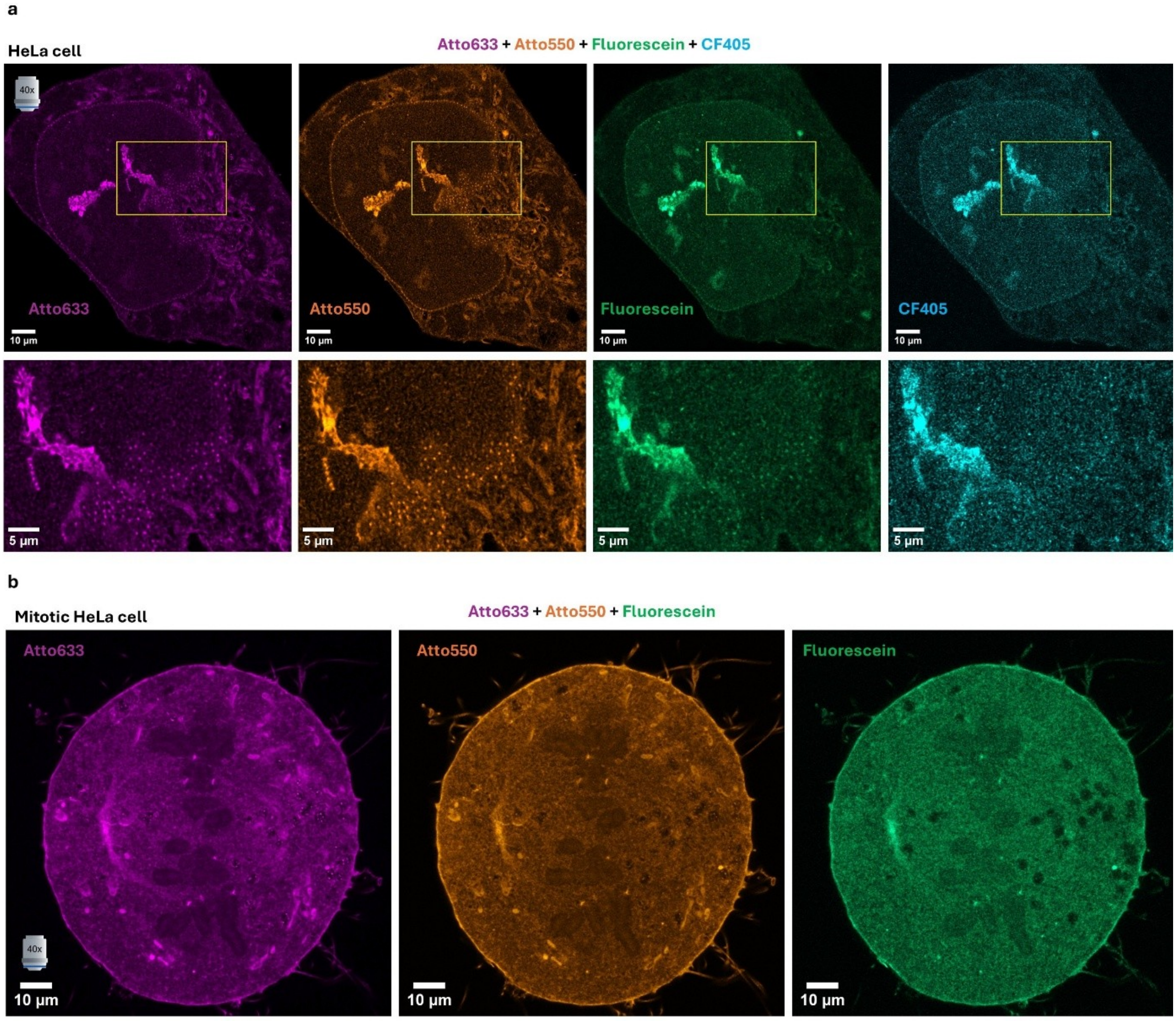


**Supplementary Figure 12. TREx-expanded cells stained with different NHS-functionalized dyes (part 2 of 4).** To evaluate if different NHS-dyes have any bias on their staining towards certain compartments of the cells, as they should have different affinities due to the variable partitioning coefficient of the dye structure (hydrophobic vs hydrophilic), we stained gels of HeLa cells with the combination of ATTO633-NHS + ATTO550-NHS + Fluorescein-NHS + CF405-NHS. **a**, HeLa cell in G0, with focus on the nuclear pores and mitochondria (yellow rectangle, zoomed below). **b**, Mitotic HeLa cell, with focus on the centrosome and the chromosomes.

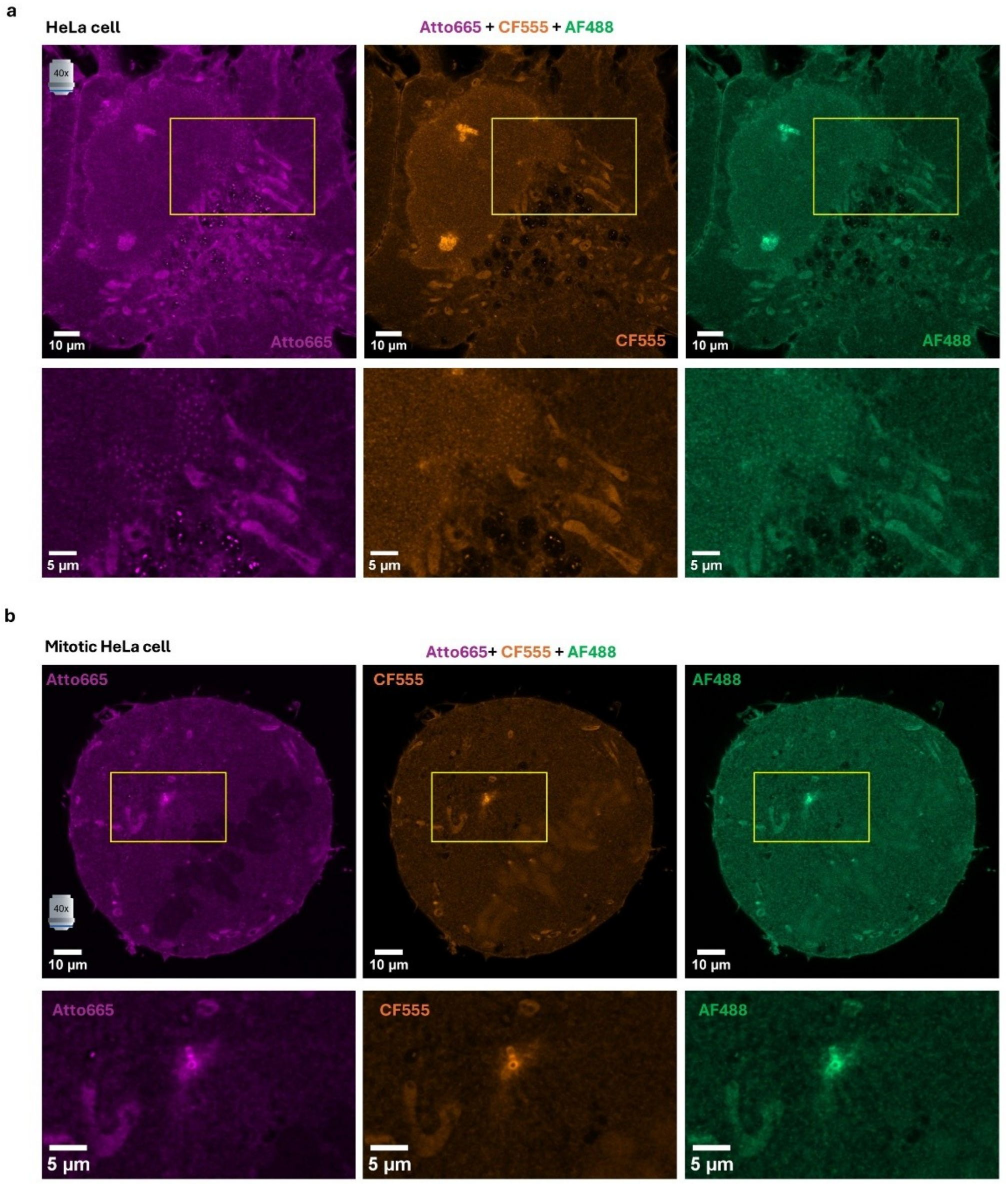


**Supplementary Fig. 13. TREx-expanded cells stained with different NHS-functionalized dyes (part 3 of 4).** To evaluate if different NHS-dyes have any bias on their staining towards certain compartments of the cells, as they should have different affinities due to the variable partitioning coefficient of the dye structure (hydrophobic vs hydrophilic), we stained gels of HeLa cells with the combination of ATTO665-NHS + CF555-NHS + AlexaFluor488-NHS. **a**, HeLa cell in G0, with focus on the nuclear pores and mitochondria (yellow rectangle, zoomed below). **b**, Mitotic HeLa cell, with focus on the centrosome (yellow rectangle, zoomed below) and the chromosomes.

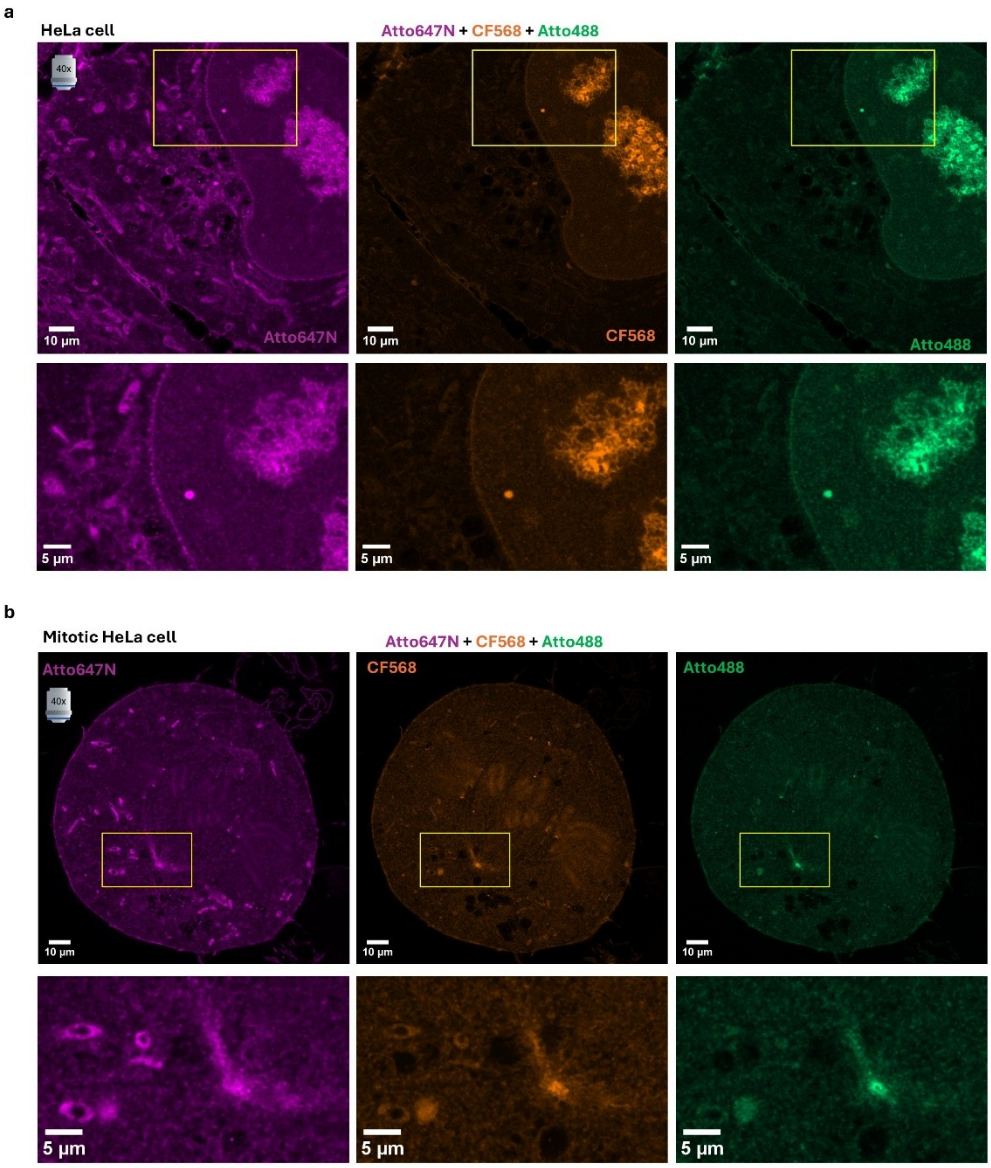


**Supplementary Fig. 14. TREx-expanded cells stained with different NHS-functionalized dyes (part 4 of 4).** To evaluate if different NHS-dyes have any bias on their staining towards certain compartments of the cells, as they should have different affinities due to the variable partitioning coefficient of the dye structure (hydrophobic vs hydrophilic), we stained gels of HeLa cells with the combination of ATTO647N-NHS + CF568-NHS + ATTO488-NHS. **a**, HeLa cell in G0, with focus on the nuclear pores and mitochondria (yellow rectangle, zoomed below). **b**, Mitotic HeLa cell, with focus on the centrosome (yellow rectangle, zoomed below) and the chromosomes.

### References


1. Baier, A., Alsheimer, M., & Benavente, R. (2007). Synaptonemal complex protein SYCP3: conserved polymerization properties among vertebrates. *Biochimica Et Biophysica Acta (BBA)-Proteins and Proteomics*, *1774*(5), 595-602.
2. Bos, P. R., Berentsen, J., Wientjes, E. Expansion microscopy resolves the thylakoid structure of spinach. *Plant Physiol.* **194**, 347-358 (2024).
3. Chang, J.B., Chen, F., Yoon, Y.G. *et al.* Iterative expansion microscopy. *Nat. Methods* **14**, 593–599 (2017).
4. Damstra, H.G.J., Mohar, B., Eddison, M., *et al.* Visualizing cellular and tissue ultrastructure using Ten-fold Robust Expansion Microscopy (TREx). *eLife* **11**, e73775 (2022).
5. Fraune, J., Schramm, S., Alsheimer, M., Benavente, R. The mammalian synaptonemal complex: protein components, assembly and role in meiotic recombination. *Exp. Cell Res.* **318,** 1340-1346 (2012).
6. Gambarotto, D., Zwettler, F.U., Le Guennec, M. *et al.* Imaging cellular ultrastructures using expansion microscopy (U-ExM). *Nat. Methods* **16**, 71–74 (2019).
7. Klimas, A., Gallagher, B.R., Wijesekara, P. *et al.* Magnify is a universal molecular anchoring strategy for expansion microscopy. *Nat. Biotechnol* .**41**, 858-869 (2023).
8. Kraft, N., Muenz, T. S., Reinhard, S., Werner, C., Sauer, M., Groh, C., & Rössler, W. (2023). Expansion microscopy in honeybee brains for high-resolution neuroanatomical analyses in social insects. *Cell and Tissue Research*, *393*(3), 489-506.
9. Kuhns, S., Schmidt, K. N., Reymann, J., Gilbert, D. F., Neuner, A., Hub, B., ... & Pereira, G. (2013). The microtubule affinity regulating kinase MARK4 promotes axoneme extension during early ciliogenesis. *Journal of Cell Biology*, *200*(4), 505-522.
10. Ku, T., Swaney, J., Park, JY. *et al.* Multiplexed and scalable super-resolution imaging of three-dimensional protein localization in size-adjustable tissues. *Nat. Biotechnol.* **34**, 973-981 (2016).
11. Louvel, V., Haase, R., Mercey, O. *et al.* iU-ExM: nanoscopy of organelles and tissues with iterative ultrastructure expansion microscopy. *Nat. Commun.* **14**, 7893 (2023).
12. M'Saad, O., Bewersdorf, J. Light microscopy of proteins in their ultrastructural context. *Nat. Commun.* **11**, 3850 (2020).
13. Sarkar, D., Kang, J., Wassie, A.T. *et al.* Revealing nanostructures in brain tissue via protein decrowding by iterative expansion microscopy. *Nat. Biomed. Eng* **6**, 1057–1073 (2022).
14. Schücker, K., Holm, T., Franke, C., Sauer, M., Benavente, R. Elucidation of synaptonemal complex organization by super-resolution imaging with isotropic resolution. *Proc. Natl. Acad. Sci. USA* **112**, 2029–2033 (2015).
15. Schmidt, K. N., Kuhns, S., Neuner, A., Hub, B., Zentgraf, H., & Pereira, G. (2012). Cep164 mediates vesicular docking to the mother centriole during early steps of ciliogenesis. *Journal of Cell Biology*, *199*(7), 1083-1101.
16. Streubel, J. M. S., Karasu, O. R., *et al.* Nek1 defines a branch of centriolar microtubule length control parallel to CP110-Cep97. *Nat. Commun.* **17**, 7330 (2026).
17. Tavakoli, M.R., Lyudchik, J., Januszewski, M. *et al.* Light-microscopy-based connectomic reconstruction of mammalian brain tissue. *Nature* **642**, 398-410 (2025).
18. Tillberg, P., Chen, F., Piatkevich, K. *et al.* Protein-retention expansion microscopy of cells and tissues labeled using standard fluorescent proteins and antibodies. *Nat. Biotechnol.* **34**, 987–992 (2016).
